# Thermo-amplifier circuit in probiotic *E. coli* for stringently temperature-controlled release of a novel antibiotic

**DOI:** 10.1101/2024.02.13.579303

**Authors:** Sourik Dey, Carsten E. Seyfert, Claudia Fink-Straube, Andreas M. Kany, Rolf Müller, Shrikrishnan Sankaran

## Abstract

Peptide drugs have seen rapid advancement in biopharmaceutical development, with over 80 candidates approved globally. Despite their therapeutic potential, the clinical translation of peptide drugs is hampered by challenges in production yields and stability. Engineered bacterial therapeutics is a unique approach being explored to overcome these issues by using bacteria to produce and deliver therapeutic compounds at the body site of use. A key advantage of this technology is the possibility to control drug delivery within the body in real time using genetic switches. However, the performance of such genetic switches suffers when used to control drugs that require post-translational modifications or are toxic to the host. In this study, these challenges were experienced when attempting to establish a thermal switch for the production of a ribosomally synthesized and post-translationally modified peptide antibiotic, darobactin, in probiotic *E. coli*. These challenges were overcome by developing a thermo-amplifier circuit that combined the thermal-switch with a T7 RNA Polymerase and its promoter that overcame limitations imposed by the host transcriptional machinery due to its orthogonality to it. This circuit enabled production of pathogen-inhibitory levels of darobactin at 40°C while maintaining leakiness below the detection limit at 37°C. More impressively, the thermo-amplifier circuit sustained production beyond the thermal induction duration. Thus, raised temperature for 2 h was sufficient for the bacteria to produce pathogen-inhibitory levels of darobactin even in the physiologically relevant simulated conditions of the intestines that include bile salts and low nutrient levels.

**Graphical Abstract:** 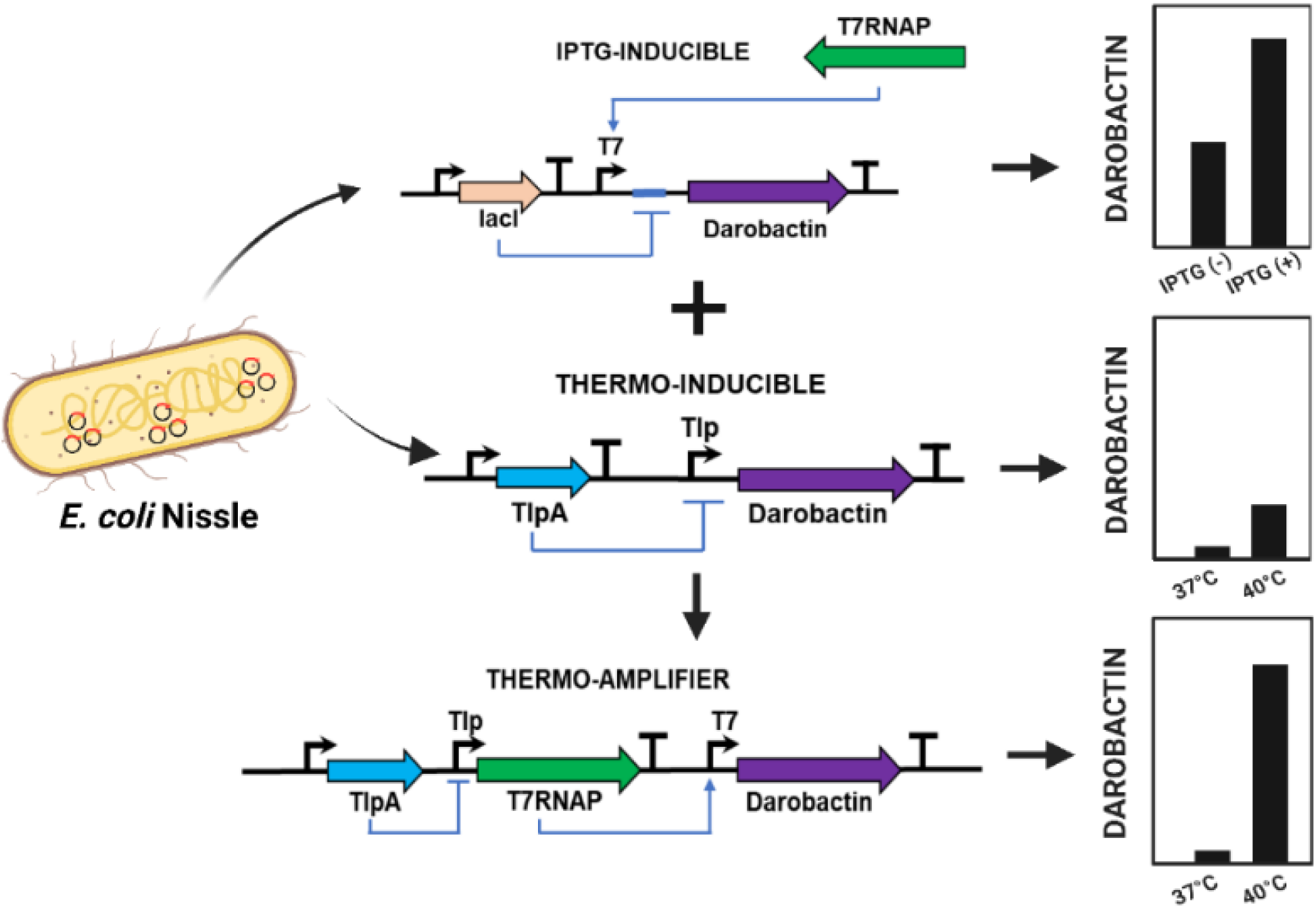

## Introduction

Among the growing spectrum of biopharmaceuticals, peptide drugs have experienced a rapid rise in development and clinical translation over the past decade with over 80 approved candidates worldwide [1,2]. These intermediate-sized drugs, ranging from 500 Da to 5 kDa, offer strong and specific interactions with their intended targets while being amenable to chemical and microbial synthesis. Furthermore, peptides can be configured into an innumerable variety of functional structures through different amino acid combinations and post-translational modifications. To discover potent candidates from this vast set of possibilities various chemical, biomolecular and computational screening techniques have been employed [3]. Accordingly, peptide drugs have been developed to act as hormones, growth factors, neurotransmitters, protease-inhibitors, immune-modulators, ion channel ligands, and anti-infective agents [4]. Nevertheless, two major hurdles exist in the translation of peptide drugs to the clinics – (i) achieving economically viable production yields, especially when their synthesis involves post-translational modifications, and (ii) limiting their instability in the body to deliver them in their active states to the disease site. A prominent example of a peptide drug facing these hurdles is darobactin, a novel antibiotic [5]. This antibiotic has shown potential in eliminating harmful Gram-negative pathogens, including highly virulent and antibiotic-resistant strains (ESKAPE classification) [5,6]. The synthesis of darobactin is possible in *E. coli* and considerable efforts are underway to improve production yields for its economically viable upscaled manufacturing [7,8,9]. As a ribosomally synthesized and post-translationally modified peptide (RiPP), the darobactin operon includes the DarA peptide followed by DarBCDE enzymes that modify it into its active form and transport it out of the cell. Recombinant production of darobactin in *E. coli* currently provides the highest yields, although this is only up to 30 mg/L over 2 days [9]. Higher yields are desired since the half-life of darobactin in the body is only 1 h, requiring the intraperitoneal administration of 50 mg darobactin per 1 kg of body weight in mice for demonstrating significant therapeutic efficacy [5].

While several strategies are underway to improve production yields and stable delivery of peptide drugs like darobactin in the body, a unique “outside-the-box” approach is the development of engineered bacterial therapeutics [10,11,12]. This emerging technology involves the genetic programming of bacteria to synthesize and administer therapeutic substances within the body. The potential applications span a wide spectrum of medical conditions including infections, inflammatory disorders and cancer [13]. These engineered bacteria can produce and deliver peptide drugs directly in the body, thereby obviating the need to externally produce these drugs in high titers. Furthermore, they can be programmed with genetic switches to control the timing and dosage of drug release, thereby ensuring drug availability only when and where it is needed. This is particularly important for antimicrobial drugs, as their indiscriminate use is known to generate antibiotic-resistant microbes [14]. While most genetic switches are activated by chemical inducers like arabinose [15] or salicylate [16] that can be taken orally, switches responsive to physical stimuli like light [17] and heat [18] offer the advantage of rapid and local activation. In particular, thermo-responsive switches are highly attractive because they can be either self-activated under fever-like conditions or externally activated using non-invasive techniques, like focused ultrasound [19]. Through these unique possibilities, engineered bacterial therapeutics can achieve on-demand production of intricate and costly biopharmaceutical drugs directly at the required body-site, thus mitigating issues associated with stability during storage, transportation, and administration. By encapsulating such bacteria in biocompatible materials, it is also possible to create smart self-replenishing drug-delivery devices to treat chronic diseases [20,21,22]. Thus, engineered bacterial therapeutics offer the possibility to provide a tailored and patient-centric treatment approach.

However, when it comes to combining genetic switches with complex biosynthetic pathways, balancing stringent control in the OFF state with sufficient production for therapeutic efficacy in the ON state is a major challenge [23]. Achieving this balance often requires considerable optimization of the genetic circuits and development of novel genetic modules [24]. In recent studies, promoter engineering enabled thermally regulated expression and release of oncolytic nanobodies [25] and tumor necrosis factor (TNF-α) [26] from the probiotic *E. coli* Nissle 1917 strain for treating solid tumors in mice models. While these studies achieved activation of bacterial gene expression within the host, the output was limited to the production and secretion of a protein without post-translational modifications. In contrast only a few studies have addressed stimuli-responsive regulation of compounds produced through enzyme cascades [27,28]. These studies often reveal that the regulated production of such compounds is more challenging compared to unmodified proteins due to greater leaky expression or lower output yields, especially when the product can be toxic to the host cell at high concentrations.

In this study, we address this challenge by engineering probiotic *E. coli* for thermo-switchable darobactin production (37 °C OFF, 40 °C ON). This approach avoids continuous release of the antibiotic from the engineered bacteria and activates it at temperatures associated with high fever. Notably, we faced the previously mentioned challenges of leaky and insufficient expression in different genetic circuits to achieve such switching. We were able to simultaneously resolve both issues by developing a genetic amplification strategy that resulted in a novel thermo-amplifier circuit with undetectable leakiness at 37°C and darobactin production levels at 40°C that inhibited the growth of the opportunistic pathogen *Pseudomonas aeruginosa* PAO1. Furthermore, we show that the thermo-amplifier circuit remains ON for several hours after only 2 h of heating, ensuring that its pathogen-killing activity can be triggered even with only a short duration of raised body temperature. Finally, the engineered strain was shown to maintain its ability to kill pathogens even in the presence of bile acids and under low-nutrient conditions that are expected to be encountered in the gastrointestinal tract. Thus, the thermo-amplifier circuit can be an efficient module to improve the performance of engineered bacterial therapeutics for the delivery of complex peptide drugs.

## Materials and Methods

### Media, bacterial strains and growth conditions

Luria-Bertani (LB) growth media used for this study was procured from Carl-Roth GmbH, Germany. Unfractionated Bovine Bile (CAS No – 8008-63-7, B3883-25G) was purchased from Merck Millipore GmbH, Germany. NEB^®^ 5-alpha Competent *E. coli* cells (C2987H) were routinely used for recombinant plasmid cloning and maintenance. The thermoamplifier constructs were assembled in chemically competent ABLE-K cells (200172) procured from Agilent Technologies (USA). ClearColi BL21 DE3 cells were obtained from BioCat GmbH (Heidelberg, Germany) and the alanine auxotrophic *Δalr Δdadx* ClearColi BL21 DE3 strain was created by Gen-H GmbH (Heidelberg, Germany) using Cre-loxP recombination technique. ClearColi were grown in LB broth supplemented with 2% (w/v) sodium chloride, with additional supplementation of 40 µg/mL D-alanine for the *Δalr Δdadx* ClearColi strain. *E. coli* Nissle 1917 (EcN) was isolated from the commercially available Mutaflor^®^ enteric coated hard capsules (Germany). *P. aeruginosa* PAO1 (DSMZ 22644) was obtained from DSMZ GmbH (Braunschweig, Germany).

### Molecular biology reagents and oligonucleotides

Q5 DNA Polymerase was used for DNA amplification and colony PCR screening (New England Biolabs GmbH, Germany). Gibson Assembly and site-directed mutagenesis was performed using the NEBuilder HiFi DNA Assembly Cloning Kit (E5520S) and KLD Enzyme Mix (M0554S) respectively. Qiagen Plasmid Kit (12125) was used for plasmid extraction and Wizard^®^ SV Gel and PCR Clean-Up System (A9282) was used for DNA purification. Oligonucleotides used in this study were synthesized from Integrated DNA Technologies (Belgium) and recombinant plasmids were verified by Sanger Sequencing (Eurofins GmbH, Germany). The sequence annotated maps of the pTlp-DarA-AT and pTAMP-DarA-AT recombinant plasmids are listed in Fig. S8 and Fig. S9. The nucleotide sequences of the genetic modules tested in this study have been listed in Table S1.

### Competent cell preparation

Electrocompetent cells of ClearColi and *Δalr Δdadx* ClearColi (alanine auxotroph) were prepared by repeated washing of bacterial pellet harvested at early exponential growth phase with 10% glycerol (v/v). Plasmids (500 ng) were transformed in the competent cells by electroporation at 1.8 kV, using 0.1 cm electroporation cuvettes (1652083) in the Bio-Rad MicropulserTM Electroporator. To prepare chemically competent EcN and EcN-T7 cells, bacterial cultures were grown overnight in LB broth at 37°C, 250 rpm and subcultured in 100 mL of fresh LB media [1:100 (v/v) dilution]. The cultures were incubated at 37°C, 250 rpm until an OD600 = 0.4 was reached. The bacteria were pelleted down by centrifugation at 4000 rpm (3363 X g), 4°C for 15 min and the supernatant was discarded. The bacterial pellet was then washed twice with ice-cold CaCl_2_ (200 mM) and once with a 1:1 mixture of CaCl_2_ (200 mM) and glycerol (10% w/v). After the final wash, the pellet was resuspended in 1 mL of CaCl_2_+glycerol mixture and stored at -80°C as 100 µL aliquots. For bacterial transformation, 1 µg of the plasmids were added to the competent cells and gently mixed by flicking. The competent cells were then chilled on ice for 30 min and transferred to a 42°C water bath for 45 s, followed by incubation on ice for 2 min. SOC media (900 μL) was added and mixed, and the cell suspension was incubated at 37°C, 250 rpm for an hour before plating on LB agar with appropriate antibiotics.

### Inducible mCherry protein production

Recombinant ClearColi BL21 DE3 strains (pNOSO-mCherry and pUC-Tlp-mCherry) were inoculated in LB-NaCl media and cultivated overnight at 30°C, 250 rpm with appropriate antibiotics. Following day, both the bacterial suspensions were subcultured in 25 mL of fresh Formulated Media (FM) [Composition:-K2HPO4 - 12.54 g/L, KH2PO4 – 2.31 g/L, D-Glucose – 4 g/L, NH4Cl – 1 g/L, Yeast Extract – 12 g/L, NaCl – 5 g/L, MgSO4 – 0.24 g/L] in a sterile glass conical flask (250 mL) at a 1:25 (v/v) ratio and incubated at 37°C, 250 rpm. Once the bacterial samples reached an OD600 of 0.4, 500 µM of IPTG was added to the pNOSO-mCherry culture, and the pUC-Tlp-mCherry culture was shifted to 40°C for further incubation. Controls were included for both strains as non-IPTG supplemented samples (pNOSO-mCherry, 37°C) and non-temperature elevated samples (pUC-Tlp-mCherry, 37°C). Post 24 h incubation, the OD600 of all the cultures was determined using the NanoDrop Microvolume UV-Vis Spectrophotometer (ThermoFisher Scientific GmbH, Germany). 1 mL of each bacterial culture was normalized to OD_600_ of 0.8 to maintain an equivalent cell count. The normalized cultures were centrifuged at 13,000 rpm (16200Xg) and washed thrice with sterile Dulbecco’s 1× PBS (Phosphate Buffer Saline) solution. 200 µL of the resuspended bacterial cultures were transferred to a clear bottom 96-well microtiter plate (Corning® 96 well clear bottom black plate, USA). The absorbance (600 nm wavelength) and mCherry fluorescence intensity (Exλ / Emλ = 587 nm/625 nm) of the respective bacterial samples were then measured in the Microplate Reader Infinite 200 Pro (Tecan Deutschland GmbH, Germany). The z-position and gain settings for recording the mCherry fluorescent intensity were set to 19442 µm and 136 respectively. Fluorescence values were normalized with the optical density of the bacterial cells to calculate the Relative Fluorescence Units (RFU) using the formula RFU = Fluorescence/OD600. For determining fold change, the following formula was used = RFU (induced culture)/RFU (uninduced culture).

### Production and purification of darobactin A

The production and purification of darobactin A was performed as described previously for darobactin D [29].

### Cultivation of the darobactin production strains

Recombinant strains were inoculated into LB (for EcN and EcN-T7 variants) and LB-NaCl (for ClearColi BL21 DE3 variants) from the glycerol stocks and cultivated overnight at 30°C, 250 rpm with appropriate antibiotics. The bacterial suspensions were then subcultured in 25 mL of fresh Formulated Media (FM) in a sterile glass conical flask (250 mL) at a 1:25 (v/v) ratio. The samples were kept at temperatures 37°C and 40°C for different temporal periods. For IPTG inducible conditions, 500 µM IPTG was added to the cultures at OD600=0.4. Bacterial growth curve was obtained by measuring OD600 at different time intervals using the Microplate Reader Infinite 200 Pro.

### Electrospray Ionization - Mass Spectrometry (ESI-MS) based darobactin quantification

For Figures 1B, 1D, 2B, 2D, 4D, 5A, 5C, S2B, S3A and S3B the following method was used: LC/ESI QTOF-MS analysis is performed on a 1260 Infinity LC in combination with a 6545A high-resolution time-of-flight mass analyzer, both from Agilent Technologies (Santa Barbara, CA, USA). Separation of 1 µl of sample is performed using a Poroshell HPH-C18 column (3.0 x 50 mm, 2.7 µm) equipped with the same guard column (3.0 x 5 mm, 2.7 µm) by a linear gradient from (A) ACN + 0.1% formic acid to (B) water + 0.1% formic acid at a flow rate of 500µl/min and a column temperature of 45°C. Gradient conditions are as follows: 0-0.5min, 95% B; 0.5-6min, 95-60.5% B; 6-9.5 min 60.5-20% B at 1500 µl/min (column cleaning), 9.5-13min 95% B down to 500µl/min (equilibrium).

**Fig. 1.**
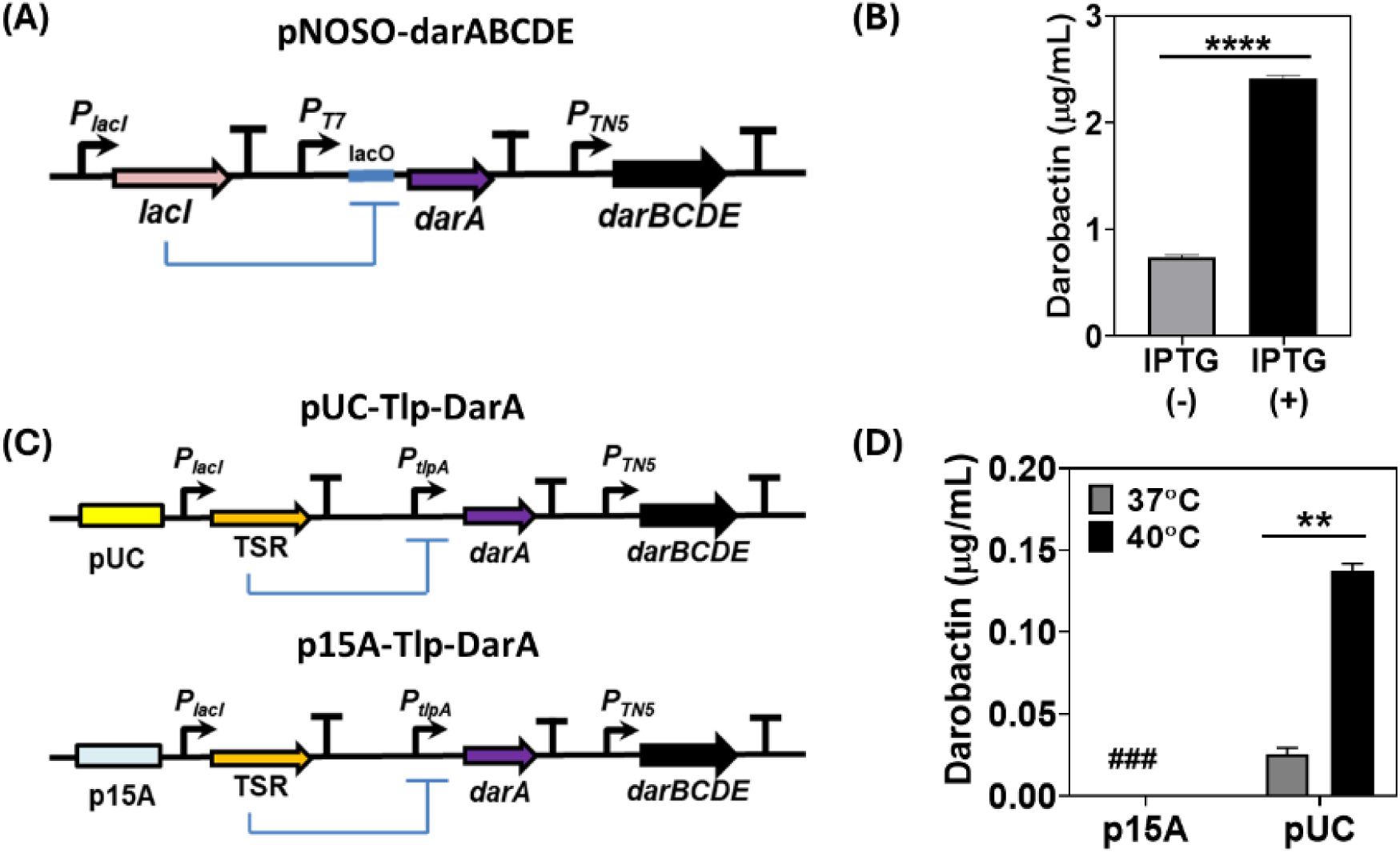
(A) Schematic representation of the pNOSO-darABCDE gene circuit (B) Darobactin concentration in the liquid medium (µg/mL) for the pNOSO-darABCDE construct in ClearColi BL21 DE3 strain after 24 h of incubation. The bacteria were induced with 500 µM of IPTG at OD_600_=0.4 in Formulated Media (FM) supplemented with 50 µg/mL kanamycin. The error bars represent standard deviation based on three independent measurements (****p *<* 0.0001 as calculated by paired t-test) (C) Schematic representation of the thermoresponsive darobactin gene circuits based on high—copy (pUC-Tlp-DarA) and medium-copy (p15A-Tlp-DarA) number replication origin. The thermosensitive repressor (TSR) represses P*_tlpA_* promoter at physiological temperature (37°C) and is derepressed at temperatures beyond >39°C (D) Darobactin concentration in the liquid medium (µg/mL) for the p15A and pUC origin based Tlp-DarA gene circuits in ClearColi BL21 DE3 strains after 24 h of incubation at 37°C and 40°C. (###) represents darobactin concentration lower than the Limit of Detection (LOD) by ESI-MS. The error bars represent standard deviation based on three independent measurements (**p = 0.0016 as calculated by paired t-test).

After separation, the LC flow enters the dual AJS ESI source set to 3500 V as capillary voltage, 40 psi nebulizer gas pressure and 7 l/min dry gas flow, and 300°C dry gas temperature. The TOF parameters used are high resolution mode (4 GHz), 140 V fragmentor and 45 V skimmer voltage. The mass spectra are acquired in the time interval of 2-6 min in full scan mode in the range m/z 150-3000 with a spectra rate of 4/s. For quantification of the darobactin, the positive charged mass [M+2H]2+ at m/z 483.7089 Da were extracted and automatically integrated using Mass Hunter software. Standards were prepared from darobactin stock solution of 100 µg/ml in mobile phase (5%A, 95%B) or blank media. Calibration was done between 0 and 1 µg/ml or 0 and 10 µg/ml with recovery rates 97-105%.

For the data in Figures 3B and 3C, the following method was used: darobactin was quantified using a Vanquish Flex UHPLC (Thermo Fisher, Dreieich, Germany), coupled to a TSQ Altis Plus mass spectrometer (Thermo Fisher, Dreieich, Germany). Samples were diluted 1:10 in PBS pH 7.4, followed by addition of 2 volumes 10%MeOH/ACN containing 15 nM diphenhydramine as internal standard. Samples were centrifuged (15 min, 4°C, 4000 rpm) before analysis and darobactin content quantified in SRM mode using a calibration curve up to 5 µM. LC conditions were as follows: column: Hypersil GOLD C-18 (1.9 μm, 100 x 2.1 mm; Thermo Fisher, Dreieich, Germany); temperature 40°C; flow rate 0.700 ml/min; solvent A: water + 0.1% formic acid; solvent B: acetonitrile + 0.1% formic acid; gradient: 0-0.2 min 10% B, 0.2-1.2 min 10-90% B, 1.2-1.6 min 90% B, 1.6-2.0 min 10% B. MS conditions were as follows: vaporizer temperature 350°C, ion transfer tube temperature 380°C, Sheath Gas 30, Aux Gas 10, Sweep Gas 2; spray voltage: 4345 V, mass transition: 483.75 – 475.083 ([M+2H]^2+^), collision energy: 11.0 V; Tube lens offset 55 V.

For analyzing experimental samples, the bacterial biomass was removed post-incubation by centrifugation [4000 rpm (3363Xg), 30 min, 4°C] and the respective supernatants were filter-sterilized with 0.22 µm filters (Carl Roth, Germany) before proceeding with ESI-MS analysis along with their respective blank controls.

### qPCR analysis for plasmid retention

Bacterial cultures were cultivated in 5 mL of LB media (supplemented with 100 μg/mL ampicillin) at 30°C with continuous shaking (250 rpm). The following day, the bacterial suspension was subcultured in 25 mL of FM both with and without antibiotic supplementation (100 µg/mL ampicillin) in a 1:25 (v/v) ratio. Bacterial cultures were immediately kept at 40°C and incubated for 24 h. Post 24 h incubation, the OD600 was measured, and fresh FM cultures were inoculated with the respective bacterial cultures at a starting OD600 of 0.01. This procedure was repeated over 5 days, resulting in ∼50 consecutive bacterial generations.

For qPCR analysis, 1 mL of the bacterial samples after every 10 generations were normalized to an OD600 of 0.8 with sterile PBS. The normalized cultures were centrifuged at 13,000 rpm (16200Xg) and washed thrice with sterile Dulbecco’s 1× PBS solution. 500 µl of the bacterial samples were then kept at 98°C for 15 min to undergo lysis in a static thermomixer (Eppendorf GmbH, Germany) according to the reported protocol [30]. The samples were then stored at -20°C for further analysis. Post collection of these bacterial samples till 50 generations, qPCR was performed using the iTaq™ Universal SYBR ^®^ Green Supermix (BioRad GmbH, Germany) in the Bio-Rad CFX96 Real time system C1000 Touch thermal cycler. To prevent sample heterogeneity, qPCR reactions were conducted with primers specific to the pUC replication origin of the recombinant plasmid. The nucleotide sequences of the qPCR primers used are listed in Table S2. The quantification curve (Cq) values were selected by the regression determination mode and the mean ΔCq was determined by the following formula: ΔCq= Cq (50^th^ generation) – Cq (10^th^ generation). The data were finally represented in the form of fold change calculated as: Mean ΔCq (With Ampicillin)/Mean ΔCq (Without Ampicillin).

### qRT-PCR analysis for gene expression

Bacterial cultures were cultivated in 5 mL of LB media (supplemented with 100 μg/mL ampicillin) at 30°C with continuous shaking (250 rpm). The following day, the bacterial suspension was subcultured in 25 mL of non-antibiotic supplemented FM in a 1:25 (v/v) ratio. Post 6 h incubation at the respective temperatures, the OD600 was determined using the NanoDrop Microvolume UV– Vis Spectrophotometer (ThermoFisher Scientific GmbH, Germany). All the bacterial cultures were normalized to OD600 of 0.8 to maintain an equivalent cell count. The normalized cultures were centrifuged at 13,000 rpm (16200Xg) and washed thrice with sterile Dulbecco’s 1× PBS solution. The bacterial pellet was then resuspended in 1X DNA/RNA protection reagent supplied with the Monarch Total RNA Miniprep Kit (NEB #T2010) (New England BioLabs GmbH, Germany) and subjected to mechanical lysis using the FastPrep-24™ 5G bead beating grinder system (MP Biomedicals Germany GmbH, Germany). Total RNA was isolated according to the manufacturer guidelines and measured at 260 nm to determine the net yield and purity. 500 ng of the total RNA from all extracted samples were then immediately converted into cDNA using the Thermo-Scientific Revertaid first strand cDNA synthesis kit (#K1622). Real time qPCR was performed using the iTaq™ Universal SYBR ^®^ Green Supermix (BioRad GmbH, Germany) in the Bio-Rad CFX96 Real time system C1000 Touch thermal cycler to determine the gene expression level of target genes. Bacterial 16S rRNA levels were also measured for all the samples as a control [31]. The specific sequences of qRT-PCR primers are listed in Table S2. The quantification curve (Cq) values were selected by the regression determination mode and the mean ΔCq values were determined by the following formula: - ΔCq= Cq (37°C) – Cq (40°C). The data were finally represented in the form of fold change calculated as 2^ΔCq^.

### *P. aeruginosa* PAO1 survival assay

*P. aeruginosa* PAO1 was inoculated into 5 mL FM from glycerol stock and subjected to overnight incubation at 37°C with 230 rpm orbital shaking. The following day, *P. aeruginosa* PAO1 was subcultured in fresh FM at 1:50 ratio (v/v) and grown till it reached log phase (OD600=0.4). The log-phase bacteria were further diluted in FM to optimize the bacterial cell number to 55X10^5^ CFU/mL. 10 µl of diluted bacterial culture were added to 100 µL of filter-sterilized supernatants (mentioned above) in a sterile 96-well U-bottom transparent plate (REF 351177, Corning, USA) to bring the final bacterial cell number to 5X10^5^ CFU/mL. The plates were incubated under static conditions for 18 h at 37°C and bacterial growth was determined by measuring absorbance at 600 nm in Microplate Reader Infinite 200 Pro.

### Darobactin production in nutrient-limited media

Bacterial cultures were cultivated overnight in 5 mL of LB media (supplemented with 100 μg/mL ampicillin) at 30°C with continuous shaking (250 rpm). The following day, the bacterial suspension was subcultured in 25 mL of FM (without antibiotic supplementation) in a 1:25 (v/v) ratio and incubated at 30°C for 24 h. Post incubation, the bacterial pellet was centrifuged at 4000 rpm (3363Xg), for 50 min, 4°C and the supernatant was discarded. The bacterial pellet was then resuspended in 25 mL of M9 Minimal Media [Na2HPO4•7H2O – 12.8 g/L, KH2PO4 – 3 g/L, NH4Cl – 1 g/L, NaCl – 0.5 g/L, D-Glucose – 4 g/L, MgSO4 – 0.24 g/L, CaCl2 – 0.011 g/L] and maintained at different incubation temperatures for 6 h. For the bile tolerance test, the bacterial pellet was resuspended in M9 Minimal Media supplemented with 0.3% (w/v) Bovine Bile [32], filter sterilized using a 50 mL Nalgene™ Rapid-Flow™ Sterile Disposable 0.22 µm Filtration Device (Catalog Number 564-0020, ThermoFisher Scientific GmbH, Germany). Post incubation, the bacterial pellet was centrifuged and discarded, whereas the filter-sterilized supernatant was further assessed for the overall darobactin concentration and its antimicrobial activity against *P. aeruginosa* PAO1.

## Results

### Functional characterization of inducible darobactin production

At first, an endotoxin-free *E. coli* strain, ClearColi BL21 DE3 [33], was engineered to produce and secrete darobactin by transforming it with the pNOSO-darABCDE plasmid [9]. In this plasmid, the DarA propeptide is encoded downstream of an IPTG-inducible T7-lacO promoter-operator complex and the DarBCDE enzyme cascade is constitutively expressed (Fig. 1A). As highlighted by Groß and coworkers before, *darBCD* encode for a permease protein (DarB), membrane fusion protein (DarC) and ATP-binding protein (DarD), all together comprising a tripartite efflux pump facilitating darobactin export from the bacterial chassis. The DarE protein is a radical *S*-adenosyl-*L-*methionine (SAM) enzyme which catalyzes the formation of the ether and C–C crosslinking in the heptapeptide core structure (W^1^–N^2^–W^3^–S^4^–K^5^–S^6^–F^7^) of the DarA propeptide [34]. This plasmid was cloned in ClearColi BL21 DE3, since this strain has been mutated to be endotoxin-free, thereby making it more immune-compatible than the regular BL21 DE3 strain that expresses the T7 RNA Polymerase (T7-RNAP) needed for recombinant gene expression. The darobactin amount secreted by this strain in Formulated Medium (FM) at 37°C following 24 h of IPTG induction (500 µM) was ∼2.5 µg/mL, detected directly in the medium by Electrospray ionization– mass spectrometry (ESI-MS). This corresponded to the previously reported minimum inhibitory concentrations (MIC) determined for the commonly tested model pathogenic strain, *P. aeruginosa* PAO1 (>2 µg/mL) [5]. However, even without IPTG induction, the system was considerably leaky and produced ∼1 µg/mL of darobactin in FM (Fig. 1B).

To achieve thermoresponsive production and secretion of darobactin, its biosynthetic gene cluster (BGC) was cloned into the thermogenetic pTlpA39-mWasabi plasmid (#Addgene 86116). In this system, a thermosensitive repressor (TSR) undergoes dimerization and represses the *PtlpA* promoter at temperatures below 39°C. Above this threshold temperature, the repressor reverts to its monomeric state allowing *PtlpA* promoter-based gene expression [18]. For the thermally activated constructs, *darA* was encoded downstream of the *PtlpA* promoter, whereas the DarBCDE enzyme cascade was constitutively expressed by the promoter *PTN5* as in the pNOSO-darABCDE plasmid. Notably, the pNOSO vector backbone has a medium-copy number p15A replicon, whereas the pTlpA39 plasmid is based on a high-copy number pMB1 replicon (pUC). Thus, two plasmids encoding thermoresponsive darobactin production modules were constructed with each of these replicons and annotated as p15A-Tlp-DarA and pUC-Tlp-DarA respectively (Fig. 1C). While no darobactin was detected in the culture medium of p15A-Tlp-DarA ClearColi strain after 24 h incubation (at both 37°C and 40°C), the ClearColi strain containing pUC-Tlp-DarA released detectable levels of darobactin in the culture medium at both incubation temperatures. The resulting darobactin concentration in the medium for the pUC-Tlp-DarA ClearColi strain at 40°C (∼0.13 µg/mL) was ∼4x fold higher than that observed at 37°C (∼0.03 µg/mL) (Fig. 1D). Although the darobactin concentrations were significantly lower than the desired MIC for inhibiting Gram-negative pathogenic strains, thermoresponsive production was confirmed. This encouraged us to optimize the performance of the gene circuit based on the high-copy number plasmid, pUC-Tlp-DarA.

To further determine the expression strength and fold-change of the inducible promoters, the *darA* was replaced by a red fluorescent reporter gene (*mCherry*) in the pNOSO and pUC-Tlp vector backbones without altering any other coding sequence (CDS) to create the pNOSO-mCherry (Fig. S1A) and pUC-Tlp-mCherry (Fig. S1B) constructs, respectively. Post IPTG induction, it was observed that mCherry production for the pNOSO-mCherry (*PT7* promoter) strain was only ∼2.5x fold higher than the uninduced control (Fig. S1C). On the other hand, the pUC-Tlp-mCherry (*PtlpA* promoter) strain had a ∼40x-fold increase in fluorescence at 40°C compared to 37°C (Fig. S1D). In addition to the superior fold change, the pUC-Tlp-DarA circuit also exhibited stronger mCherry production [in terms of relative fluorescence unit (RFU)] at 40°C (Fig. S1F) than the IPTG induced pNOSO-mCherry circuit (Fig. S1E). However, this contrasting trend between the darobactin and mCherry production by the same pUC-Tlp based recombinant plasmids suggested that the genetic circuits encoding darobactin production might be influenced by alternate factors like the *darA:darE* transcript ratio [8] or intracellular toxicity [29], rather than the production rate of DarA alone. In accordance with our observations regarding pNOSO-darABCDE, there was considerable basal level expression of mCherry observed for pNOSO-mCherry samples grown without any IPTG supplementation. On the other hand, mCherry expression was tightly regulated by the TSR protein in pUC-Tlp-mCherry samples at 37°C in line with our previous report [35]. This shows that TSR protein-based gene regulation is more effective than the conventional lacI repressor protein which is known for its inability to mediate strong repression [36].

Another major factor that distinguishes darobactin production from that of mCherry is that darobactin could exert a detrimental effect on the growth of the bacterial host [5,9]. To validate this hypothesis, constitutive expression of darobactin was tested in ClearColi by exchanging the thermoresponsive elements in pUC-Tlp-DarA with the *PT7* promoter (pT7-DarA), placed directly upstream of the d*arA* gene for driving its constitutive transcription by the cognate T7-RNAP genomically encoded in ClearColi. As expected, this caused a significant reduction in the bacterial biomass (∼100 mg per 25 mL of liquid culture) in comparison to the IPTG-induced pNOSO-darABCDE construct (∼430 mg per 25 mL of liquid culture) when cultivated at 37°C for 24 h (Fig. S2A). A considerable reduction in the overall darobactin concentration (∼0.09 µg/mL) was also observed for the pT7-DarA ClearColi strain when compared to the IPTG-induced pNOSO-darABCDE ClearColi strain (Fig. S2B). It could be deduced that the constitutive expression of darobactin by the pT7-DarA construct imposed a significant metabolic burden on the producer strain. Therefore, inducible expression of darobactin was essential to achieve higher darobactin concentration without compromising the metabolic fitness of the microbial chassis.

### Selection of an efficient antibiotic supplementation-free plasmid retention strategy

Before further optimization of thermoresponsive darobactin production, strategies to retain the pUC-Tlp-DarA plasmid in an antibiotic-free manner were tested. Earlier reports on engineered probiotics have highlighted the importance of maintaining the genetic stability of the heterologous protein-encoding genes during bacterial growth [37]. Although genetic stability can be achieved by integrating the heterologous genes in the host genome, studies have shown that this could significantly reduce the overall production [38] and secretion [39] levels of recombinant proteins. As we were unable to observe darobactin production in the liquid cultures using a medium-copy number replication origin (p15A ori), we decided not to proceed with genomic integration of the recombinant genetic modules for this study. As low but detectable levels of darobactin were observed for the thermoresponsive genetic circuit based on the high-copy number pUC replication origin, we chose this recombinant construct for assessing the two commonly employed plasmid retention strategies based on nutrient auxotrophy [40] and toxin-antitoxin dependent post-segregational killing [41]. For auxotrophy based plasmid retention, the alanine racemase genes (*alr* and *dad*X) essential for D-alanine biosynthesis in ClearColi were deleted from the bacterial genome. The resultant double knockout Δ*alr* Δ*dadX* ClearColi strain could therefore only be cultivated when the growth medium was supplemented with D-alanine. The *alr* gene with its native promoter was then amplified from *E. coli* DH5α genomic DNA and cloned into the vector backbone of pUC-Tlp-DarA to create the pTlp-DarA-alr construct (Fig. 2A). Once transformed with this recombinant plasmid, the constitutive expression of the *alr* gene could then sustain the growth of the Δ*alr* Δ*dadX* ClearColi strain even in the absence of D-alanine supplementation and support plasmid retention. For the toxin-antitoxin (TA) based plasmid retention strategy, a type II TA gene pair, *txe-axe*, originally isolated from *Enterococcus faecium* was used [42]. Once the Txe endoribonuclease protein is expressed, it is actively bound by its less stable counterpart, the Axe antitoxin protein, thereby preventing it from targeting the intracellular RNA population. However, if the recombinant strain loses the *txe-axe* gene-harboring plasmid, a drastic decline in Axe antitoxin concentration releases the Txe endoribonuclease, causing spontaneous RNA cleavage and bacterial growth arrest [43]. Recent studies have shown that incorporating the *txe-axe* TA module in the recombinant constructs allowed stable plasmid retention in *E. coli* over several generations, both under *in-vitro* [44] and *in-vivo* conditions [45]. The *txe-axe* gene pair was amplified from the pUC-GFP-AT (Addgene #133306) plasmid and cloned into pUC-Tlp-DarA to create the pTlp-DarA-AT plasmid (Fig. 2C). The antibiotic supplementation-free retention plasmids, pTlp-DarA-alr and pTlp-DarA-AT, were transformed in Δ*alr* Δ*dadX* ClearColi and unmodified ClearColi strains, respectively and cultivated in FM for 24 h, both with and without antibiotic supplementation (100 µg/mL ampicillin). Post incubation, no significant difference of darobactin concentrations (liquid medium) was observed for the strains containing either pTlp-DarA-alr (Fig. 2B) or pTlp-DarA-AT (Fig. 2D) in the absence of the antibiotic when compared to in the presence of it. The fold change (∼4x) in darobactin production in response to the incubation temperatures also remained unaffected during this period. These results suggested that both approaches could effectively retain the engineered plasmids during bacterial growth and simultaneous darobactin production for 24 h even in the absence of antibiotic supplementation. As no distinct advantage of either strategy was noticed in terms of darobactin production in the tested temporal period, preference was given to the *txe-axe* TA system for its ability to be used across several related *E. coli* strains without them having to undergo any permanent genomic modifications [25].

**Fig. 2.**
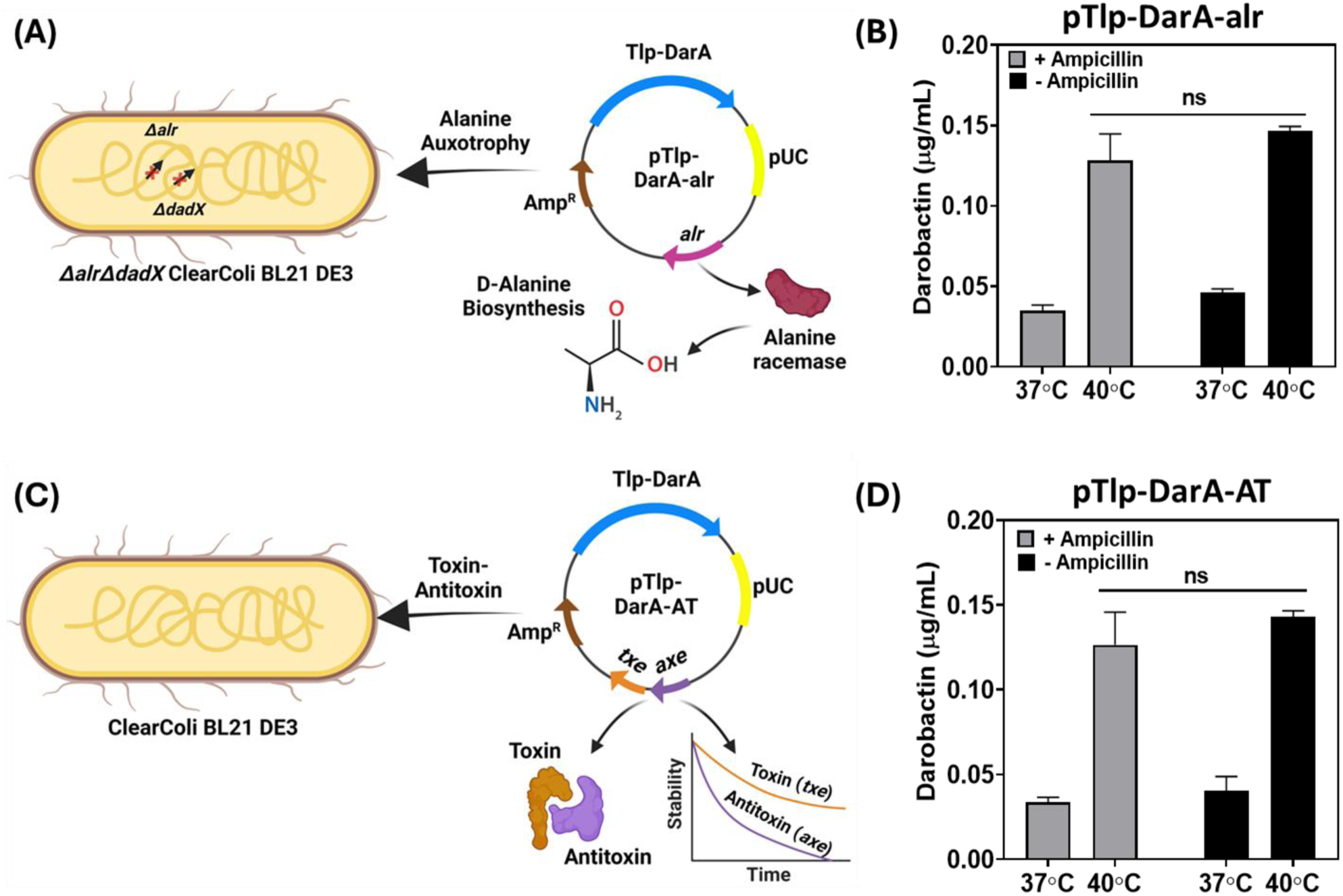
(A) Schematic representation of the pTlp-DarA-alr construct transformed in Δ*alr* Δ*dad*X ClearColi BL21 DE3 strain. The integration of the alanine racemase (*alr*) gene in the pUC-Tlp-DarA vector backbone facilitates alanine racemase-based conversion of L-alanine to D-alanine, responsible for bacterial cell wall synthesis in the auxotrophic strain (B) Darobactin concentration (µg/mL) in the liquid medium for the pTlp-DarA-alr construct after 24 h of incubation at 37°C and 40°C, both with and without ampicillin (100 µg/mL) supplementation. The error bars represent the standard deviation based on three independent measurements (^ns^p = 0.1482 as calculated by paired t-test) (C) Schematic representation of the pTlp-DarA-AT construct transformed in ClearColi BL21 DE3 strain. The integration of the *txe* toxin - *axe* antitoxin (TA) gene pair in the pUC-Tlp-DarA vector backbone mediate plasmid retention by its endoribonuclease activity and differential protein stability rates (D) Darobactin concentration (µg/mL) in the liquid medium for the pTlp-DarA-AT construct after 24 h of incubation at 37°C and 40°C, both with and without ampicillin (100 µg/mL) supplementation. The error bars represent the standard deviation based on three independent measurements (^ns^p = 0.2584 as calculated by paired t-test).

### Optimized genetic circuit to increase darobactin production

From the previous results, it was determined that the *PT7* promoter with the cognate T7-RNAP was able to support high levels of darobactin production but poor inducible fold-changes. On the other hand, the *PtlpA* promoter along with its TSR was found to mediate thermoresponsive darobactin production with an appreciable fold change between 37°C and 40°C but poor overall concentration in the liquid medium. Therefore, to obtain a higher darobactin concentration while maintaining thermoresponsive production, we decided to combine components of both inducible systems to design a thermo-amplifier circuit (pTAMP). In this genetic circuit, *darA* gene expression is mediated by the *PT7* promoter and the production of its cognate T7-RNAP [amplified from the mBP-T7-RNAP plasmid (Addgene #74096)] is controlled by the TSR regulated *PtlpA* promoter (Fig. 3A). Notably, such a circuit could not be tested in ClearColi, since the constitutively expressed *t7-RNAP* gene integrated into the bacterial genome would interfere with the performance of the recombinant circuit. Thus, as an alternative, we chose the probiotic EcN strain for testing the performance of the recombinant plasmid because of its prior use as a living therapeutic agent [46]. However, before proceeding with the testing of the pTAMP circuit we needed to assess whether darobactin production could be sustained by the EcN strain. For that, we first transformed the pNOSO-darABCDE plasmid in the EcN-T7 strain (DSMZ 115365), an engineered EcN variant with a genomic integration of the constitutively expressed *t7-RNAP* gene [47]. Post IPTG induction, the darobactin levels obtained from this recombinant EcN-T7 strain after 24 h in FM was over 3x-fold higher (∼8.15 µg/mL) than that from the ClearColi strain. This suggested that *PT7* driven production of darobactin could also be sustained in EcN without compromising the metabolic fitness of the microbial chassis. Furthermore, similar to what was observed with the pNOSO-darABCDE ClearColi strain, the corresponding EcN-T7 strain also exhibited high basal level production of darobactin (∼4.7 µg/mL) even in the absence of IPTG (Fig. S3A). We also wanted to verify whether the unmodified EcN strain was able to produce darobactin when transformed with the thermoresponsive pTlp-DarA-AT plasmid. Post 24 h incubation, the darobactin concentration in the liquid medium (FM) without antibiotic supplementation for the pTlp-DarA-AT EcN strain was found to be ∼ 0.02 µg/mL and ∼0.07 µg/mL at 37°C and 40°C, respectively (Fig. S3B). This showed that the darobactin concentrations from EcN are very similar to that of the ClearColi strain, with an overall fold change of ∼3x at higher temperature (40°C). Both these experiments validated that EcN was a suitable microbial chassis for constructing and testing the thermo-amplifier circuit.

**Fig. 3.**
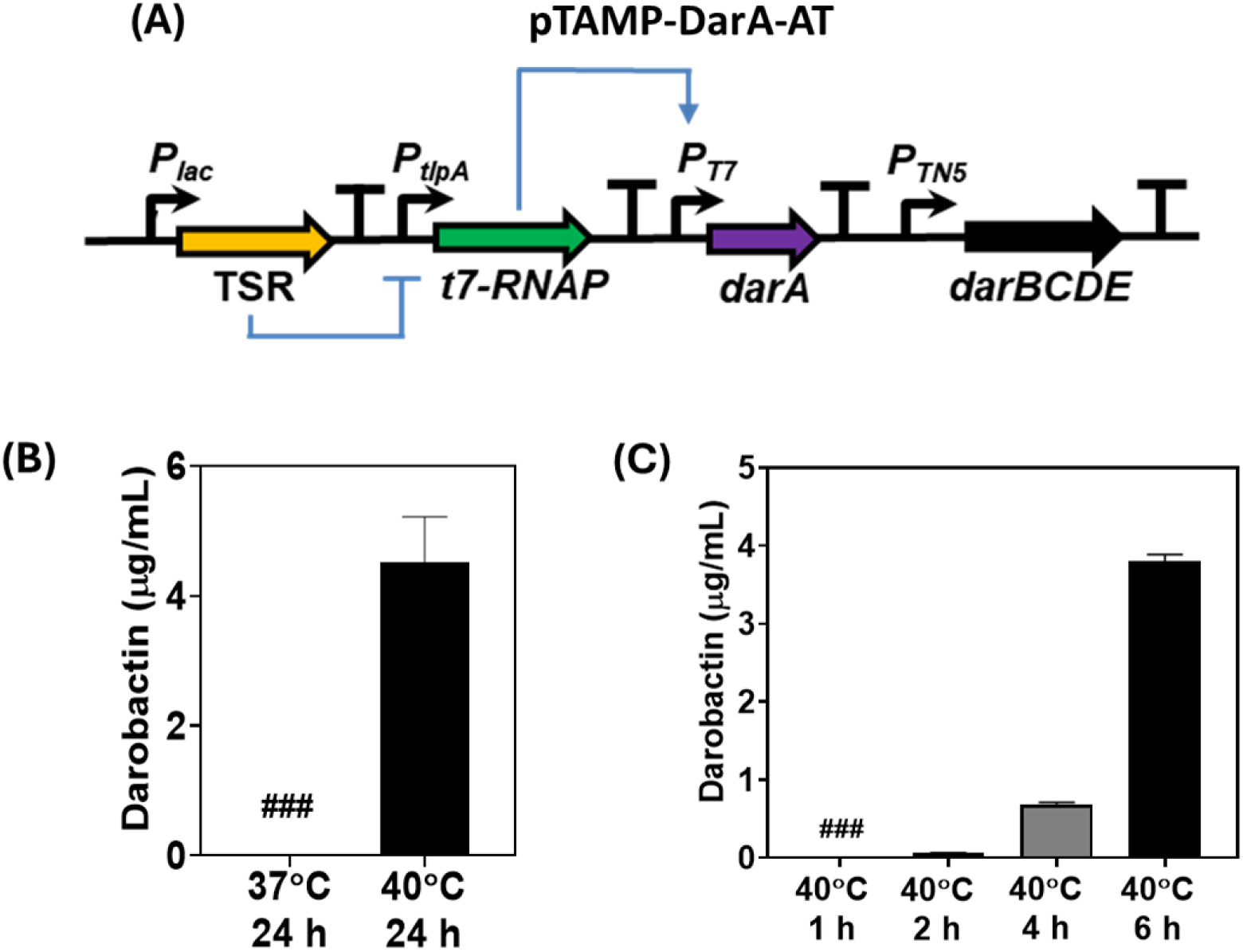
(A) Schematic representation of the pTAMP-DarA-AT genetic circuit. The TSR represses the *t7-RNAP* gene at incubation temperatures <37°C, beyond which the TSR de-repression leads to *t7-RNAP* expression and *P_T7_* promoter based darobactin production (B) Darobactin concentration in the liquid medium (µg/mL) for the pTAMP-DarA-AT construct in *E. coli* Nissle 1917 (EcN) after 24 h of incubation at 37°C and 40° C respectively. The error bars represent standard deviation based on three independent measurements. (###) represents darobactin concentrations lower than the Limit of Detection (LOD) by ESI-MS (C) Darobactin concentration in the liquid medium (µg/mL) for the pTAMP-DarA-AT EcN strain after 1, 2, 4 and 6 h of incubation at 40° C, respectively. Error bars represent standard deviation from three independent measurements. (###) represents darobactin concentrations lower than the LOD by ESI-MS.

Our key hypothesis was that post thermal induction (40°C), the thermo-amplifier circuit should be able to offer higher levels of darobactin production, while showing minimal darobactin expression at physiological temperature (37°C). Surprisingly, when the thermo-amplifier circuit was constructed in a plasmid, pTAMP-DarA-AT and tested in EcN, drastic improvement in the darobactin concentration (liquid medium) was observed at 40°C (∼4.5 µg/mL), while its presence was below the detection limit of 0.01 µg/mL at 37°C after 24 h incubation in FM (Fig. 3B). It is to be also noted that the darobactin concentration achieved with the pTAMP-DarA-AT EcN strain was ∼1.6x-fold higher than that obtained by the pNOSO-darABCDE ClearColi strain post-IPTG induction. In addition to darobactin production, we also wanted to assess whether the *txe-axe* TA module could mediate recombinant plasmid retention over several bacterial generations without external antibiotic supplementation. To do so, the pTAMP-DarA-AT EcN strain was cultivated for ∼50 consecutive generations in FM at 40°C over a period of 5 days both with and without ampicillin supplementation (100 µg/mL). qPCR analysis of bacterial populations after 10 and 50 generations revealed no significant difference between the mean ΔCq fold change for the samples cultivated without antibiotic supplementation when compared to the antibiotic supplemented samples (Fig. S4). This suggested that the *txe-axe* TA system ensured that the microbial chassis retained the plasmid at similar copy numbers even after ∼50 generations of cell division in the absence of external selection pressure.

Since darobactin is responsible for the growth inhibition of Gram-negative bacteria, including *E. coli*, the effect of its overproduction in the EcN strain in terms of the bacterial growth kinetics needed to be tested. While the growth rate of the pTAMP-DarA-AT EcN strain was slightly reduced compared to the pTlp-DarA-AT EcN and wild type EcN strains, at both 37°C (Fig. S5A) and 40°C (Fig. S5B) incubation temperatures, there was no significant temporal delay observed for the strains to reach their maximal cell density. The growth curves also showed that all the strains had an exponential increase in their cell density up to 6 h, corresponding to their log phase beyond which slower growth rates were observed. This motivated us to determine whether thermal induction for shorter periods of time could produce detectable levels of darobactin in the liquid medium. Following this, we cultivated the pTAMP-DarA-AT EcN strain in FM for 1, 2, 4 and 6 h (periods corresponding to the log phase) at both the incubation temperatures of 37°C and 40°C. None of the bacterial samples incubated at 37°C showed detectable levels of darobactin in the liquid medium. However, for the bacterial samples incubated at 40°C, detectable levels of darobactin could be observed beyond 2 h incubation (∼0.06 µg/mL), with darobactin concentrations reaching ∼0.7 µg/mL and ∼3.8 µg/mL upon 4 h and 6 h incubation, respectively (Fig. 3C). High darobactin levels in the liquid medium after 6 h incubation ascertained that the majority of darobactin was produced during the exponential growth phase of the recombinant strain, beyond which both the darobactin concentration and bacterial growth rate underwent gradual stagnation.

To gain a better understanding, we conducted rt-PCR analysis for the *t7-RNAP* and *darA* genes of the pTAMP-DarA-AT EcN strain, post-incubation at 37°C and 40°C for 6 h. rt-PCR analysis revealed that the gene expression levels of *t7-RNAP* and *darA* were ∼9 and ∼19x fold higher at 40°C than at 37°C, respectively (Fig. 4A & Fig. 4B). In comparison, no significant upregulation in the *PtlpA* promoter driven *darA* gene expression could be observed for the pTlp-DarA-AT EcN strain at 40°C (11.6 ± 0.3 cycles) compared to 37°C (11.5 ±1.7 cycles) after 6 h incubation in FM (Fig. S6). This could explain the overall low darobactin concentration observed for the *PtlpA* promoter-driven darobactin production by the pTlp-DarA-AT EcN strain after thermal induction. Furthermore, the quantification cycle (Cq) value of *darA* gene at 40°C for the pTAMP-DarA-AT construct (EcN) was 8.1 ± 1.2 cycles, which formed the logical basis for obtaining a higher darobactin concentration compared to the pTlp-DarA-AT construct (EcN) after a 6 h incubation period. This analysis suggested that the lower production levels of pTlp-DarA are due to the producer strains’ inability to effectively transcribe the gene rather than the potential toxicity of DarA. This could be due to one or more of the variety of mechanisms involving transcription factors and the RNA polymerase that mediate transcription initiation [48], elongation [49] and termination [50]. However, the limiting mechanism doesn’t seem to affect transcription driven by the phage-derived T7-RNAP.

**Fig. 4.**
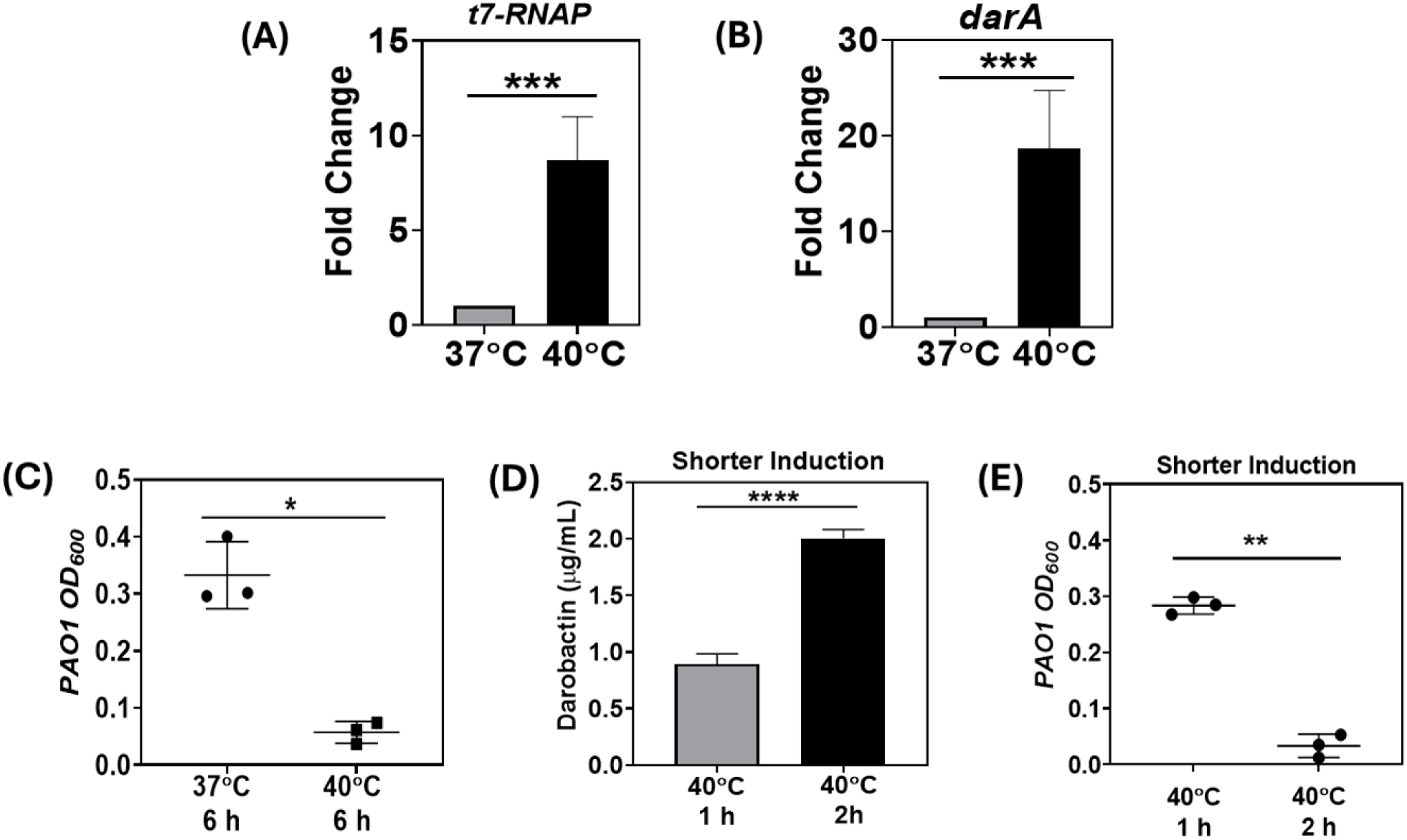
(A) Fold change in *t7-RNAP* gene expression driven by the *P_tlpA_* promoter of pTAMP-DarA-AT construct in EcN at 37°C and 40°C after 6 h incubation in FM (without antibiotic supplementation). The error bars represent standard deviation based on six independent measurements (***p =0.0004 as calculated by paired t-test) (B) Fold change in *darA* gene expression driven by the *P_T7_* promoter of pTAMP-DarA-AT construct in EcN at 37°C and 40°C after 6 h incubation in FM (without antibiotic supplementation). The error bars represent standard deviation based on six independent measurements (***p =0.0009 as calculated by paired t-test) (C) End-point absorbance (OD_600_) of *P. aeruginosa* PAO1 after 18 h incubation at 37°C in filter-sterilized FM previously sustaining the growth of pTAMP-DarA-AT EcN strain. The FM collected from the 40°C incubated samples demonstrate significant growth inhibition of *P. aeruginosa* PAO1. The error bars represent standard deviation based on three independent measurements (*p = 0.0253 as calculated by paired t-test) (D) Darobactin concentration in the liquid medium (µg/mL) for the pTAMP-DarA-AT EcN strain grown in FM (without antibiotic supplementation) after 1 h and 2 h of shorter thermal induction (40°C), followed by an additional 5 h and 4 h of incubation at 37°C, respectively. The error bars represent standard deviation based on three independent measurements (****p <0.0001 as calculated by paired t-test) (E) End-point absorbance (OD_600_) of *P. aeruginosa* PAO1 after 18 h incubation at 37°C in filter-sterilized FM previously sustaining the growth of pTAMP-DarA-AT EcN strain. The FM collected from the 2 h thermally induced (40°C) samples followed by an additional incubation at 37°C for 4 h demonstrate significant growth inhibition of *P. aeruginosa* PAO1. The error bars represent standard deviation based on three independent measurements (**p = 0.0014 as calculated by paired t-test).

Next, we determined whether the high levels of darobactin produced by the thermo-amplifier genetic circuit imparted antimicrobial activity against Gram-negative pathogens. We selected the commonly used laboratory strain, *P. aeruginosa* PAO1, as the indicator pathogen for our darobactin-based antimicrobial screening assays. Although *P. aeruginosa* is not a common enteric pathogen, it has been reported to colonize the gastrointestinal tract and cause gut-derived sepsis, Shanghai fever, and severe respiratory infections, especially in infants and immunocompromised patients [40]. MIC experiments with pure darobactin revealed that concentrations above ∼2 µg/mL could inhibit the growth of *P. aeruginosa* PAO1 in FM-based liquid cultures (Fig. S7), in accordance with previous studies [9]. Encouraged by these results, we tested whether the darobactin produced by the pTAMP-DarA-AT EcN strain could exert an antimicrobial activity against *P. aeruginosa* PAO1. Filter-sterilized supernatants collected from the pTAMP-Dar-AT EcN cultures after 6 h incubation at 37°C and 40°C, were inoculated with the *P. aeruginosa* PAO1 strain. Post overnight incubation, only the bacterial supernatants collected from the 40°C incubated pTAMP-Dar-AT EcN cultures inhibited the growth of *P. aeruginosa* PAO1. In contrast, the bacterial supernatants collected from their counterparts incubated at 37°C failed to inhibit the growth of *P. aeruginosa* PAO1 (Fig. 4C). Although this was an interesting observation, thermal activation by high fever (>39 °C) or focused ultrasound cannot be expected to last for 6 h. For instance, high fever triggered by bacterial pyrogens either do not last longer than a couple hours [51] or are brought down by medication [52]. Therefore, thermal induction for the darobactin production cascade for a prolonged period of 6 h might not be most suitable for treating pathogen. Since the T7-RNAP produced by the thermo-amplifier circuit via thermal induction should remain active even after the temperature is lowered, we expected darobactin production to be sustained for longer. Therefore, we decided to test the minimal thermal induction threshold that would be suitable for both activation and sustained production of darobactin from the recombinant EcN strain. For that, we subjected the pTAMP-DarA-AT EcN strain to thermal induction at 40°C for a 1 h and 2 h period, and then lowered the temperature to 37°C for the following 5 h and 4 h, respectively. After the 6 h incubation period, ∼0.9 µg/mL and ∼2 µg/mL of darobactin could be detected in the liquid medium for the cultures subjected to the short thermal induction periods of 1 h and 2 h, respectively (Fig. 4D). The higher darobactin concentration observed for the 2 h induced samples were also able to effectively inhibit the growth of *P. aeruginosa* PAO1 (Fig. 4E), whereas the 1 h induced samples did not demonstrate significant antimicrobial activity. Considering the results in Fig. 3C, which show that 2 h of thermal induction doesn’t produce more than 0.6 µg/mL, the results of Fig. 4E suggests that the darobactin production is sustained beyond the duration of thermal induction. Thus, thermal induction at 40°C for 2 h followed by an additional 4 h incubation at 37°C was sufficient to activate and sustain darobactin production from the pTAMP-DarA-AT EcN strain to inhibit *P. aeruginosa* PAO1 growth.

When considering potential applicability in the gastrointestinal tract, the recombinant pTAMP-DarA-AT EcN strain must be able to perform under conditions of nutrient limitation and the presence of bile acids [53]. Therefore, we tested our engineered EcN strains in a nutrient-limited medium resembling the gut luminal content [54]. The pTAMP-DarA-AT EcN strain was inoculated in M9 Minimal Media and subjected to thermal induction for 2 h at 40°C, with an additional incubation of 4 h at 37°C. However, this medium was not able to support the growth of sufficient biomass to generate pathogen-inhibiting quantities of darobactin concentration (<0.1 µg/mL). Thus, we mimicked the high-biomass delivery modality used for probiotics and living therapeutics by first growing the bacteria to a sufficient biomass in FM medium at 30 °C (OFF state), then testing their darobactin production capacity in M9 Minimal Medium at 40 °C (ON state). The harvested biomass from FM medium was re-suspended in M9 Minimal Media and subjected to thermal induction at 40°C for 2 h, followed by an additional incubation of 4 h at 37°C. Post thermal induction, darobactin concentration in the supernatants reached up to ∼1.8 µg/mL, whereas the control samples incubated at 37°C for 6 h produced only ∼0.3 µg/mL of darobactin (Fig. 5A). Pre-cultivating the bacterial biomass for 24 h may have resulted in the basal level expression of T7-RNAP, leading to the leaky darobactin production from the resuspended samples incubated at 37°C. However, the ∼7x fold higher concentration of darobactin in the supernatants collected from the thermally induced bacterial samples demonstrated significant growth inhibition of *P. aeruginosa* PAO1 compared to the control samples (Fig. 5B). Finally, we tested the darobactin production capacity of the pTAMP-DarA-AT EcN strain in the presence of bile stress, which is known to promote antibiotic resistance [55] and biofilm formation [56] in *P. aeruginosa* PAO1. For this, the pre-cultivated biomass of the pTAMP-DarA-AT EcN strain was resuspended in M9 Minimal Media supplemented with 0.3% (w/v) Bovine Bile, and then subjected to a 2 h thermal induction at 40°C followed by an additional incubation at 37°C for 4 h. Upon further analysis, ∼2.2 µg/mL of darobactin could be observed in the supernatants of the thermally induced bacterial samples, whereas ∼0.5 µg/mL of darobactin could be detected in the supernatants of the control samples incubated at 37°C (Fig. 5C). The similar darobactin concentration fold change (∼5x) suggested that bile stress did not significantly affect the darobactin production capacity of the thermo-amplifier EcN strain. In addition, the supernatants of the thermally induced bacterial samples also showed significant growth inhibition of *P. aeruginosa* PAO1 compared to the control samples suggesting that bile supplementation did not reduce the antimicrobial activity of darobactin (Fig. 5D). These experiments confirmed that the thermo-amplifier EcN strain could sustain darobactin production in the presence of stress factors, such as bile and nutrient limitations.

**Fig. 5.**
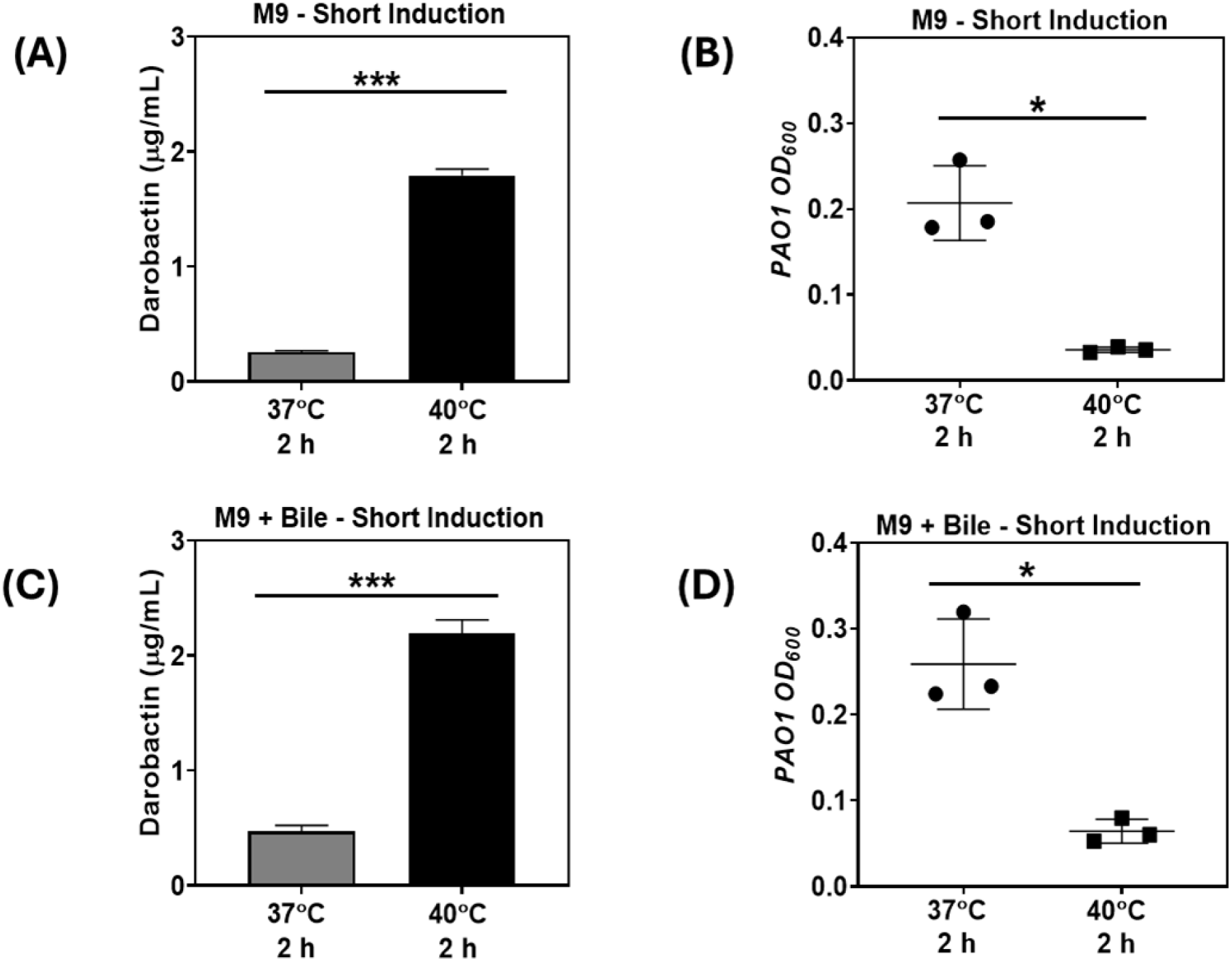
(A) Darobactin concentration (μg/mL) in M9 Minimal Media sustaining the pre-cultivated biomass of pTAMP-DarA-AT EcN strain (no antibiotic supplementation) after 2 h incubation at 37°C and 40°C (thermal induction), followed by an additional 4 h incubation at 37°C. The error bars represent standard deviation based on three independent measurements (***p=0.0004 as calculated by paired t-test) (B) End-point absorbance (OD_600_) of *P. aeruginosa* PAO1 after 18 h incubation at 37°C in filter-sterilized M9 Minimal Media containing the pTAMP-DarA-AT EcN strain. The samples subjected to thermal induction (40°C) for 2 h followed by an additional incubation at 37°C for 4 h demonstrate significant growth inhibition of *P. aeruginosa* PAO1. The error bars represent standard deviation based on three independent measurements (*p = 0.0234 as calculated by paired t-test) (C) Darobactin concentration (μg/mL) in M9 Minimal Media supplemented with 0.3% (w/v) Bovine Bile sustaining the pre-cultivated biomass of pTAMP-DarA-AT EcN strain (no antibiotic supplementation) after 2 h incubation at 37°C and 40°C (thermal induction), followed by an additional 4 h incubation at 37°C. The error bars represent standard deviation based on three independent measurements (***p=0.0005 as calculated by paired t-test) (D) End-point absorbance (OD_600_) of *P. aeruginosa* PAO1 after 18 h incubation at 37°C in filter-sterilized M9 Minimal Media supplemented with 0.3% (w/v) Bovine Bile, containing the pTAMP-DarA-AT EcN strain. The samples subjected to thermal induction (40°C) for 2 h followed by an additional incubation at 37°C for 4 h demonstrate significant growth inhibition of *P. aeruginosa* PAO1. The error bars represent standard deviation based on three independent measurements (*p = 0.0328 as calculated by paired t-test).

## Discussions

In this study, we have demonstrated the challenges in regulating the production of an enzymatically synthesized novel antibiotic and developed a solution to overcome those challenges. Specifically, we show how one inducible system (IPTG) results in high production of darobactin but also suffers from significant basal level expression. This basal level expression can occur due to the affinity of the lacI repressor protein to related auto-inducer molecules, like lactose (disaccharide) and galactose (monosaccharide) present in the growth media as reported previously [57]. Such cross-reactivity to simple carbohydrates cannot be reasonably avoided when pathogen inhibition is to be tested under therapeutically relevant conditions. On the other hand, such cross-reactivity does not occur in the TlpA-based thermoresponsive genetic switch [58] but its transcriptional limitations can restrict heterologous protein production in the microbial chassis and reduce their application potential. This was observed in our case, where the standard thermal induction circuit could not reach the expected darobactin production levels in the liquid medium that was observed for the IPTG inducible system. By combining components from both systems, we developed a thermo-amplifier circuit that achieved undetectable levels of leaky production and high levels of thermally induced darobactin production in the growth medium. This was done by exploiting the strong transcriptional rate of the T7 promoter as previously reported for the opto-T7-RNAP and related systems [59,60]. The orthogonality of the T7 RNA Polymerase system also seems to overcome limitations imposed by the host transcription machinery, which in the case of *PtlpA*-driven darobactin production resulted in very low yields. Thus, for the first time, we have demonstrated the advantages of combining thermo-genetics with the prolific strength and orthogonality of the T7-RNAP-T7 promoter combination and demonstrated the superior performance of this circuit for darobactin production. Furthermore, a favorable consequence of the thermo-amplifier circuit design is the sustained production of darobactin even after only a short duration (2 h) of thermal induction. Such short durations of raised body temperature can more conceivably be achieved by high fever or focused ultrasound. This performance was retained even in the presence of bile salts and under low-nutrient conditions, which better mimic the environment in the intestines [53]. Previous reports have shown considerable improvements in engineering *E.* coli-based strains, to sense and eliminate *P. aeruginosa* PAO1, both under *in-vitro* [61] and *in-vivo* [62] conditions. The engineered strains could detect low levels of the quorum sensing molecule, N-acyl homoserine lactone (3OC12-HSL) and undergo auto-lysis to release anti-*P. aeruginosa* toxin, pyocin S5 and reduce gut infections in animal models. As pointed out by Hwang and co-workers engineered EcN strains could colonize the mice intestine for a period of 3 weeks, which provided an extended temporal window to facilitate complete pathogen clearance. Furthermore, EcN strains have been engineered to produce surface-displayed adhesins or curli fiber matrices that improve their retention and local density in the intestines for drug delivery [63,64]. These promising engineering strategies can be combined with our thermo-amplifier darobactin circuit to improve its applicability in the body.

In the current study, all these components are encoded in a single plasmid for ease of manipulation and testing across different *E. coli* strains. The plasmid hosts a toxin-antitoxin system that ensures retention and desired darobactin production levels without antibiotic supplementation. Nevertheless, the presence of genes encoding for darobactin exporter enzymes in the recombinant plasmid also increased the darobactin tolerance of the EcN as shown by the MIC values (>16 µg/mL compared to 1.6 µg/mL of wild type EcN) without affecting the growth, which is consistent with the results previously observed for a standard *E. coli* BL21 DE3 strain [9]. This raises concerns regarding horizontal transfer of the plasmids leading to darobactin resistance in other microbes. In follow-up studies, we will explore two strategies to mitigate this potential risk – (i) encode the darBCD cluster in the genome, and (ii) encapsulate the bacteria in polymeric matrices that prevent horizontal gene transfer [65]. The second approach, based on engineered living materials, also offers additional benefits conferred by the material component such as protection of the bacteria in the body and their physical biocontainment [22]. This strategy is being increasingly explored to overcome challenges of balancing efficacy and biosafety in engineered living therapeutics [20,66].

While the current study has demonstrated the effectiveness of the darobactin-producing thermo-amplifier circuit using a model pathogen, *P. aeruginosa* PAO1, this drug has broad spectrum antimicrobial activity against several Gram-negative pathogens [5,9]. The applicability of this system to treat other infections caused by pathogens like *Klebsiella*, *Salmonella*, *Shigella*, etc. is of great interest for future studies. Beyond darobactin, the thermo-amplifier circuit could also be adapted to achieve highly regulated production of other enzymatically synthesized therapeutic compounds. For example, it can be applied to numerous bioactive RiPPs like darobactin whose BGCs have been recently elucidated and reconstructed for heterologous expression in *E. coli* [67]. Finally, beyond thermal induction, the amplification strategy using the T7 RNA Polymerase and its cognate promoter could be adapted for other inducible systems that suffer from poor performance either due to high leaky expression, low post-induction expression levels, or interference from host transcriptional machinery.

## Conclusions

Living therapeutics represents an exciting frontier for realizing long-term, low-cost and controlled drug release within the body. However, creating leak-free and rapidly responsive genetic switches to control drug release remains a major challenge hindering the field, especially for drugs that require post-translational modification and are toxic to the production host. In this study, we have presented a novel strategy to achieve stringent genetic control over the production and release of a complex post-translationally modified peptide drug, darobactin, using probiotic bacteria. While well-known IPTG- and thermo-responsive genetic switches achieved either leaky or insufficient production, the careful combination of parts from both switches resulted in a thermo-amplifier circuit that overcame both issues. The production and release of darobactin at pathogen-inhibitory levels using an antibiotic-free plasmid retention system under physiologically relevant conditions highlights the possibility of developing such probiotics as delivery vehicles for drugs that are challenging to mass-produce at low cost.

## Declarations

## Acknowledgements

We would like to thank Mikhail Shapiro for the pTlpA39-mWasabi plasmid (Addgene #86116), Chris Barnes for the pUC-GFP AT plasmid (Addgene #133306) and Paul Freemont for the mBP-T7-RNAP plasmid (Addgene #74096). We thank Nicole Frankenberg-Dinke for providing us with the EcN-T7 strain (DSMZ 115365). We thank Sanjana Balaji Kuttae for her help with the construction of the recombinant plasmids. We thank Ha Rimbach-Nguyen for assisting with the ESI-MS analysis of darobactin concentration in bacterial supernatants. All the schematic figures were generated using Biorender.

## Ethics approval and consent to participate

Not applicable.

## Consent for publication

Not applicable.

## Availability of data and materials

The authors declare that the data supporting the findings of this study are available within the paper and its Supplementary Information files. Should any raw data files be needed in another format they are available from the corresponding author upon reasonable request. Data are located in controlled access data storage at the INM - Leibniz Institute for New Materials.

## Author Contributions

S.D. designed and constructed the plasmids, performed experiments, and analyzed the data. C.E.S. provided the purified darobactin standard. C.F.S. and A.M.K. performed the quantitative estimation of darobactin in the extracellular media. S.S. and R.M. conceived and oversaw the overall project. S.D. and S.S. wrote the manuscript. C.E.S. and R.M. revised the manuscript. All authors read and approved the manuscript.

## Supplementary Information

Fig. S1: Comparative mCherry production by the pNOSO-mCherry and pUC-Tlp-mCherry plasmids in ClearColi BL21 DE3; Fig. S2: Comparative analysis of biomass and darobactin production by the pNOSO-darABCDE and pT7-DarA plasmids in ClearColi BL21 DE3; Fig. S3: Comparative analysis of darobactin production by the pNOSO-darABCDE and pTlp-DarA-AT plasmids in EcN-T7 and EcN strains respectively; Fig. S4: qPCR analysis-based assessment of pTAMP-DarA-AT recombinant plasmid retention in EcN with and without antibiotic supplementation; Fig. S5: Growth kinetics of the engineered strains at 37°C and 40°C; Fig. S6: *darA* gene expression level by the pTlp-DarA-AT plasmid; Fig. S7: Minimum Inhibitory Concentration (MIC) of darobactin for *Pseudomonas aeruginosa* PAO1 strain; Fig. S8 & Fig. S9: Sequence annotated maps of pTlp-DarA-AT and pTAMP-DarA-AT recombinant plasmids; Table S1: Nucleotide sequences of genetic modules; Table S2: Primer sequences for qRT-PCR and qPCR analysis.

