## Supplementary information for "Thermo-amplifier circuit in probiotic *E. coli* for stringently temperature-controlled release of a novel antibiotic"

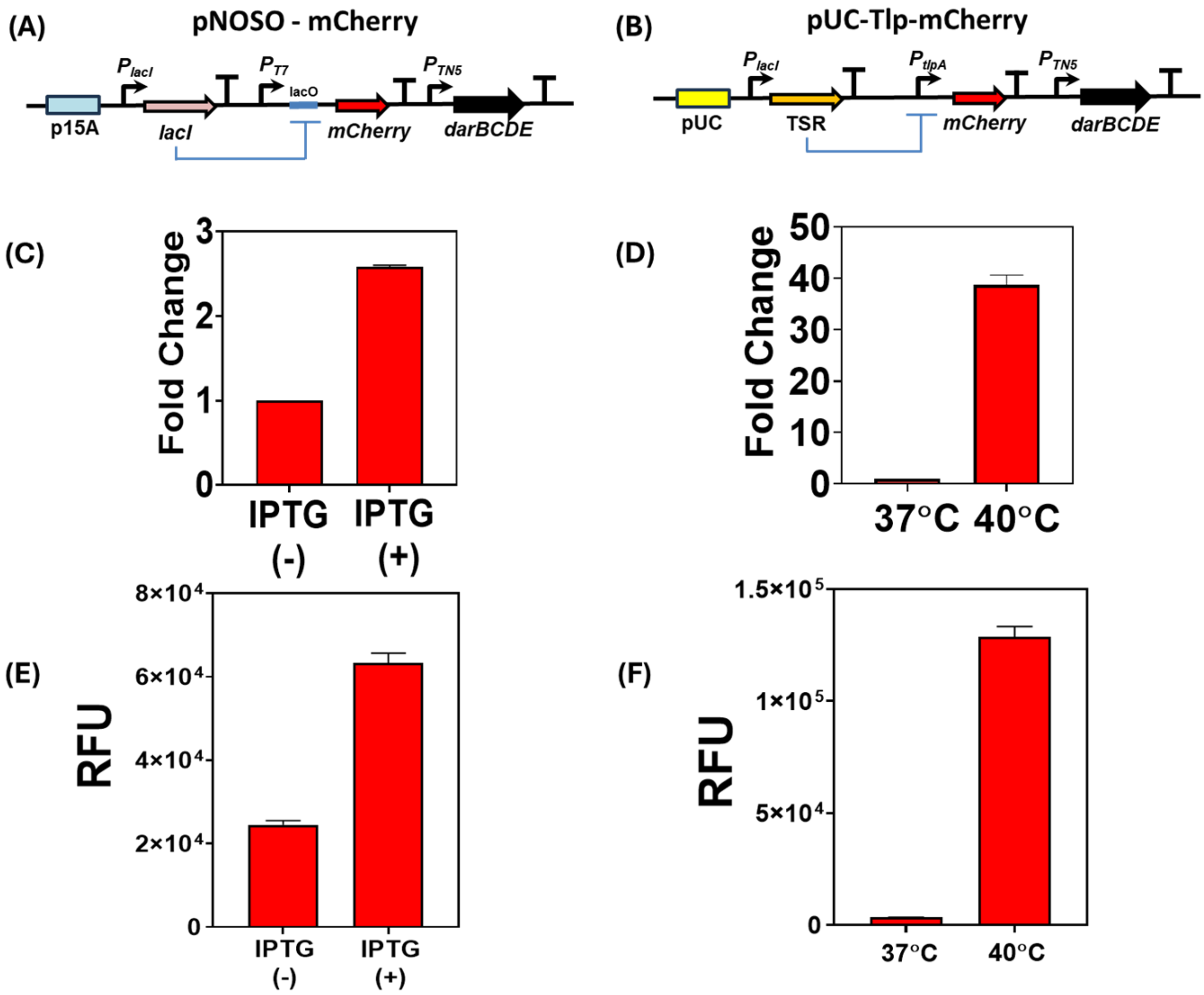


**Supplemental Figure 1.** (A) Schematic representation of the pNOSO-mCherry genetic circuit (B) Schematic representation of the pUC-Tlp-mCherry genetic circuit (C) Fold Change of $P_{T7}$ promoter driven mCherry expression in pNOSO-mCherry ClearColi strain after 24 h incubation at 37°C. The data represents the increase in mCherry expression post 500 µM IPTG induction [IPTG (+)] in comparison to the non-induced sample [IPTG (-)]. The error bars represent the standard deviation based on three independent measurements (D) Fold Change of $P_{tlpA}$ promoter driven mCherry expression in pUC-Tlp-mCherry ClearColi strain after 24 h incubation at 37°C and 40°C respectively. The data represents the increase in mCherry expression at 40°C incubation temperature in comparison to the 37°C incubated samples. The error bars represent the standard deviation based on three independent measurements (E) Relative Fluorescence Units (RFU) of mCherry produced by the pNOSO-mCherry ClearColi strain after 24 h incubation at 37°C. The data represents the increase in mCherry expression post 500 µM IPTG induction [IPTG (+)] in comparison to the non-induced sample [IPTG (-)]. The error bars represent the standard deviation based on three independent measurements (F) Relative Fluorescence Units (RFU) of mCherry produced by the pUC-Tlp-mCherry ClearColi strain after 24 h incubation at 37°C and 40°C respectively. The data represents the increase in mCherry expression when incubated at 40°C in comparison to 37°C. The error bars represent the standard deviation based on three independent measurements.

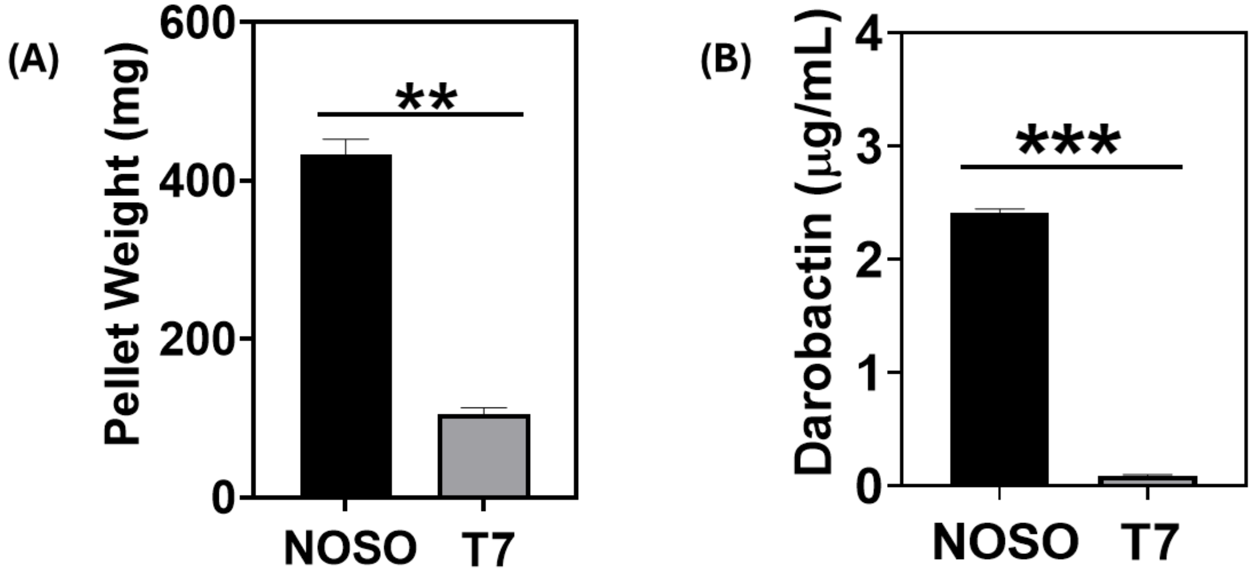


**Supplemental Figure 2.** (A) Biomass comparison in terms of the bacterial pellet wet weight [in milligrams (mg)] obtained from a 25 mL culture after 24 h incubation at 37°C. The pNOSO-darABCDE ClearColi strain was induced with 500 µM IPTG at log phase ($OD_{600}$=0.4), whereas the pT7-DarA ClearColi strain was not subjected to IPTG supplementation. The error bars represent standard deviation based on three independent measurements (**p < 0.001 as calculated by paired t-test) (B) Darobactin concentration (in µg/mL) in the liquid medium of IPTG-induced pNOSO-darABCDE ClearColi strain in comparison to the non-IPTG supplemented pT7-DarA ClearColi strain after 24 h incubation at 37°C. The error bars represent standard deviation based on three independent measurements (***p < 0.0001 as calculated by paired t-test).

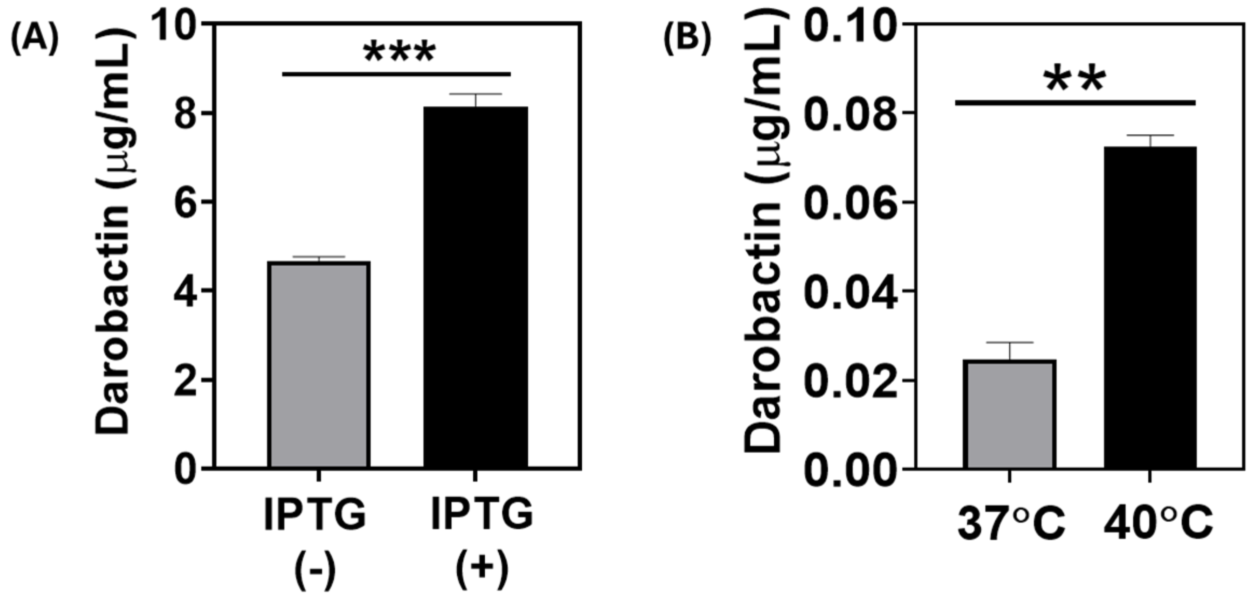

**Supplemental Figure 3.** (A) Darobactin concentration in the liquid medium (in µg/mL) of the pNOSO-darABCDE *E. coli* Nissle 1917 – T7 (EcN-T7) strain after 24 h of incubation at 37°C. For IPTG induction, the bacterial sample was grown till $OD_{600}$ = 0.4 in Formulated Media (FM) supplemented with 50 µg/mL kanamycin and then induced with 500 µM of IPTG. The error bars represent standard deviation based on three independent measurements (***p = 0.001 as calculated by paired t-test) (B) Darobactin concentration in the liquid medium (in µg/mL) of the pTlp-DarA-AT *E. coli* Nissle 1917 (EcN) strain after 24 h incubation at 37°C and 40°C respectively (without antibiotic supplementation). The error bars represent standard deviation based on three independent measurements (**p = 0.0058 as calculated by paired t-test).

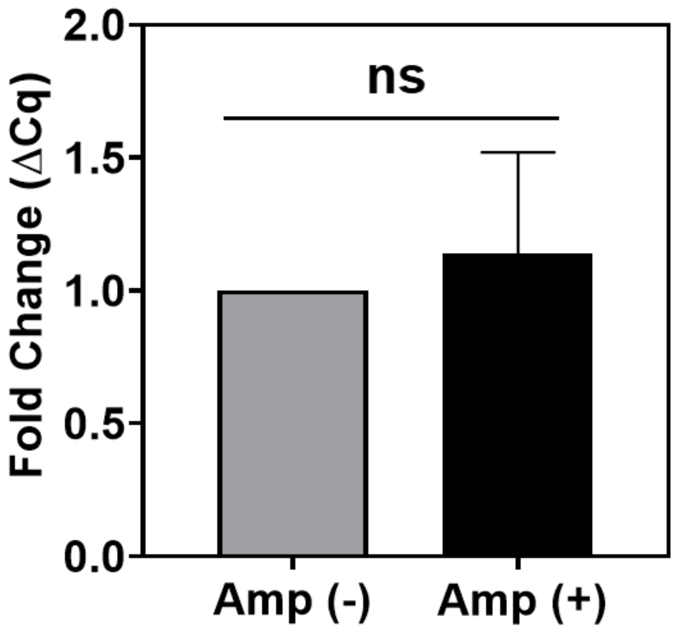


**Supplemental Figure 4.** qPCR analysis-based detection of the pTAMP-DarA-AT recombinant plasmid in EcN cultivated with and without ampicillin (100 µg/mL) supplementation at 40°C, compared over 10 and 50 generation numbers. The error bars represent standard deviation based on three independent measurements ($^{ns}$p = 0.5955 as calculated by paired t-test).

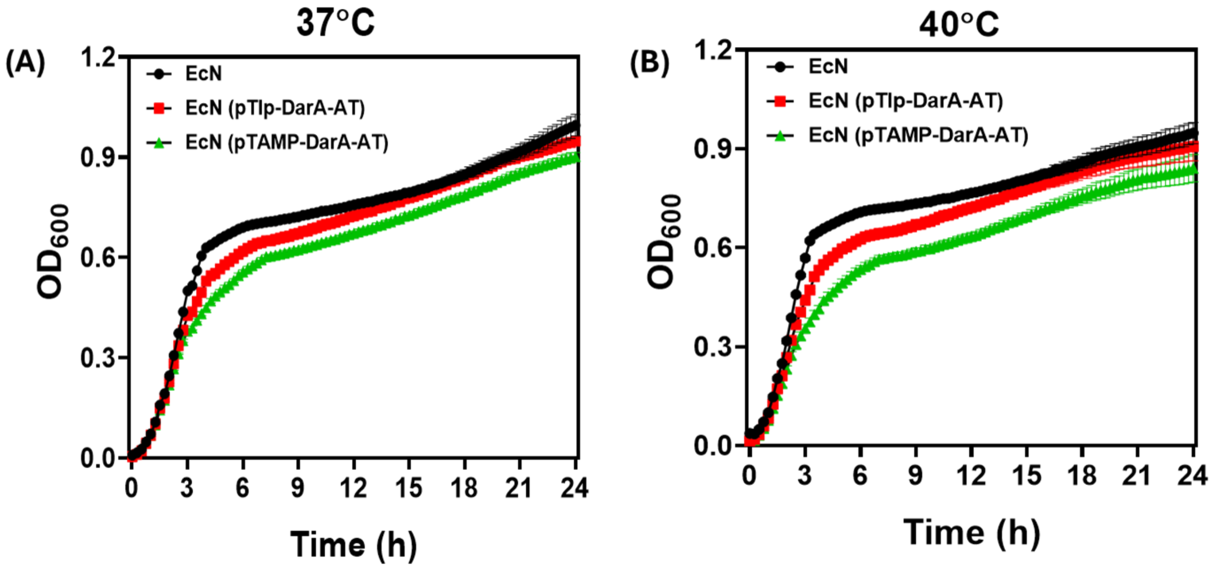


**Supplemental Figure 5.** (A) Growth Kinetics of Wild Type *E. coli* Nissle (EcN), thermo-responsive *E. coli* Nissle (pTlp-DarA-AT EcN) and thermo-amplifier *E. coli* Nissle (pTAMP-DarA-AT EcN) strains at 37°C for a 24 h incubation period. The samples were shaken continuously, and their Optical Density ($OD_{600}$) measured at equal time intervals. The error bars represent standard deviation based on three independent measurements (B) Growth Kinetics of Wild Type *E. coli* Nissle (EcN), thermo-responsive *E. coli* Nissle (pTlp-DarA-AT EcN) and thermo-amplifier *E. coli* Nissle (pTAMP-DarA-AT EcN) strains at 40°C for a 24 h incubation period. The samples were shaken continuously, and their Optical Density ($OD_{600}$) measured at equal time intervals. The error bars represent standard deviation based on three independent measurements.

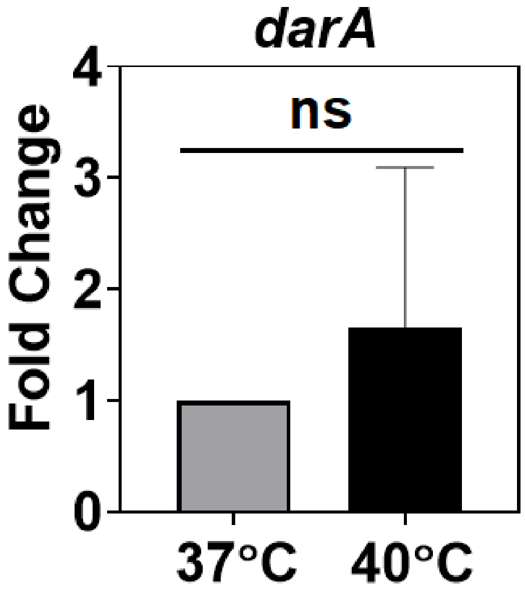

**Supplemental Figure 6.** Fold change in *darA* gene expression driven by the $P_{tlpA}$ promoter of pTlp-DarA-AT construct in EcN at 37°C and 40°C after 6 h incubation in FM (without antibiotic supplementation). The error bars represent standard deviation based on six independent measurements ($^{ns}p$ = 0.3137 as calculated by paired t-test).

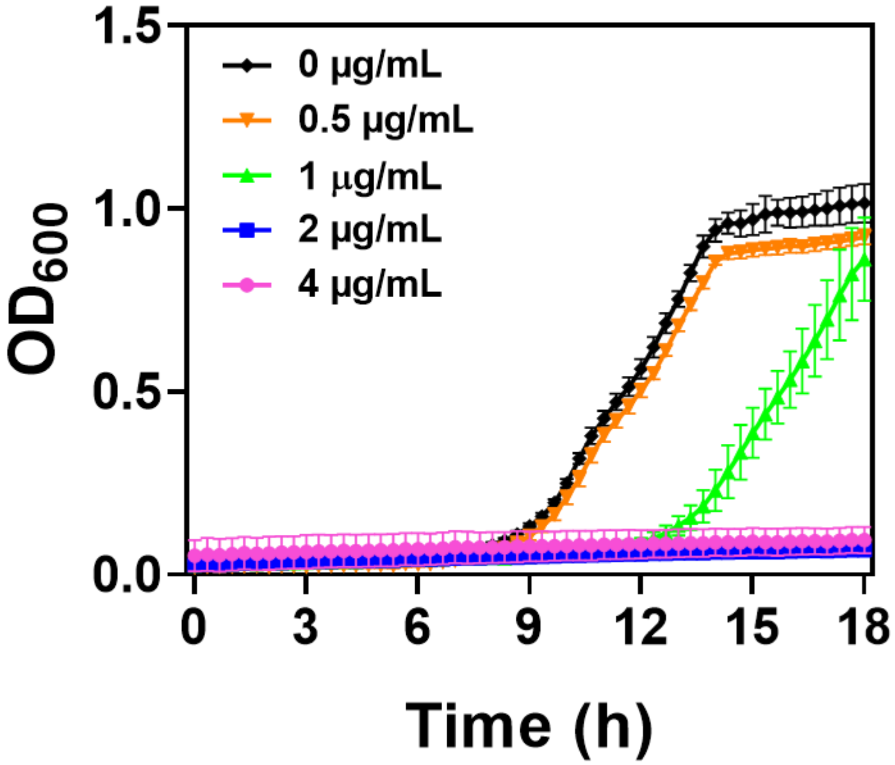


**Supplemental Figure 7.** Growth Kinetics of *Pseudomonas aeruginosa* PAO1 (DSMZ 22644) in Formulated Media (FM) supplemented with 0.5, 1, 2 and 4 µg/mL of darobactin respectively. Non-darobactin supplemented samples were used as a control (0 µg/mL). Absorbance at 600 nm (Optical Density or OD600) of the samples was recorded at regular intervals at 37°C incubation temperature along with continuous orbital shaking. The error bars represent standard deviation based on three independent measurements.

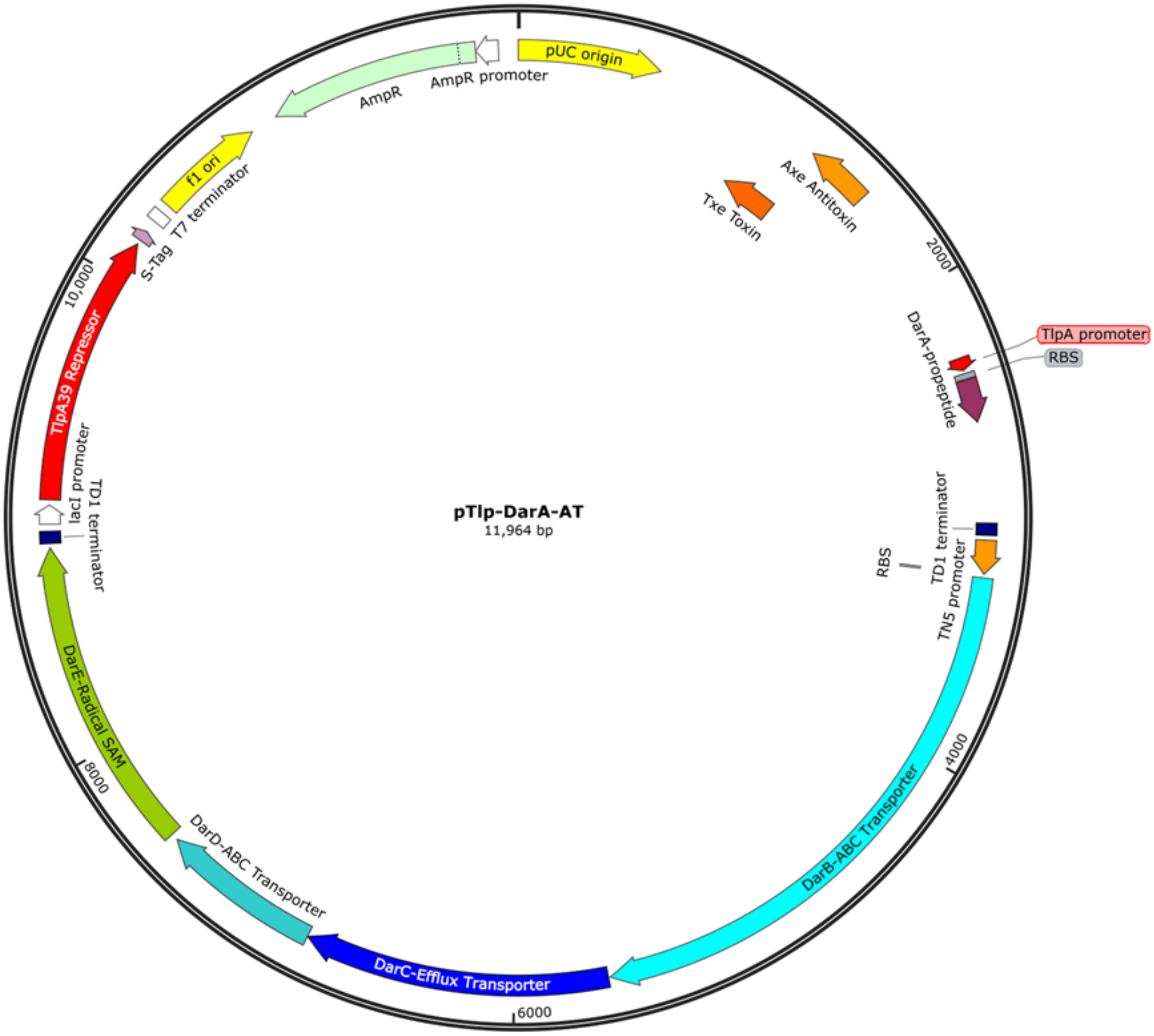


**Supplemental Figure 8.** Sequence annotated map of the pTlp-DarA-AT recombinant plasmid

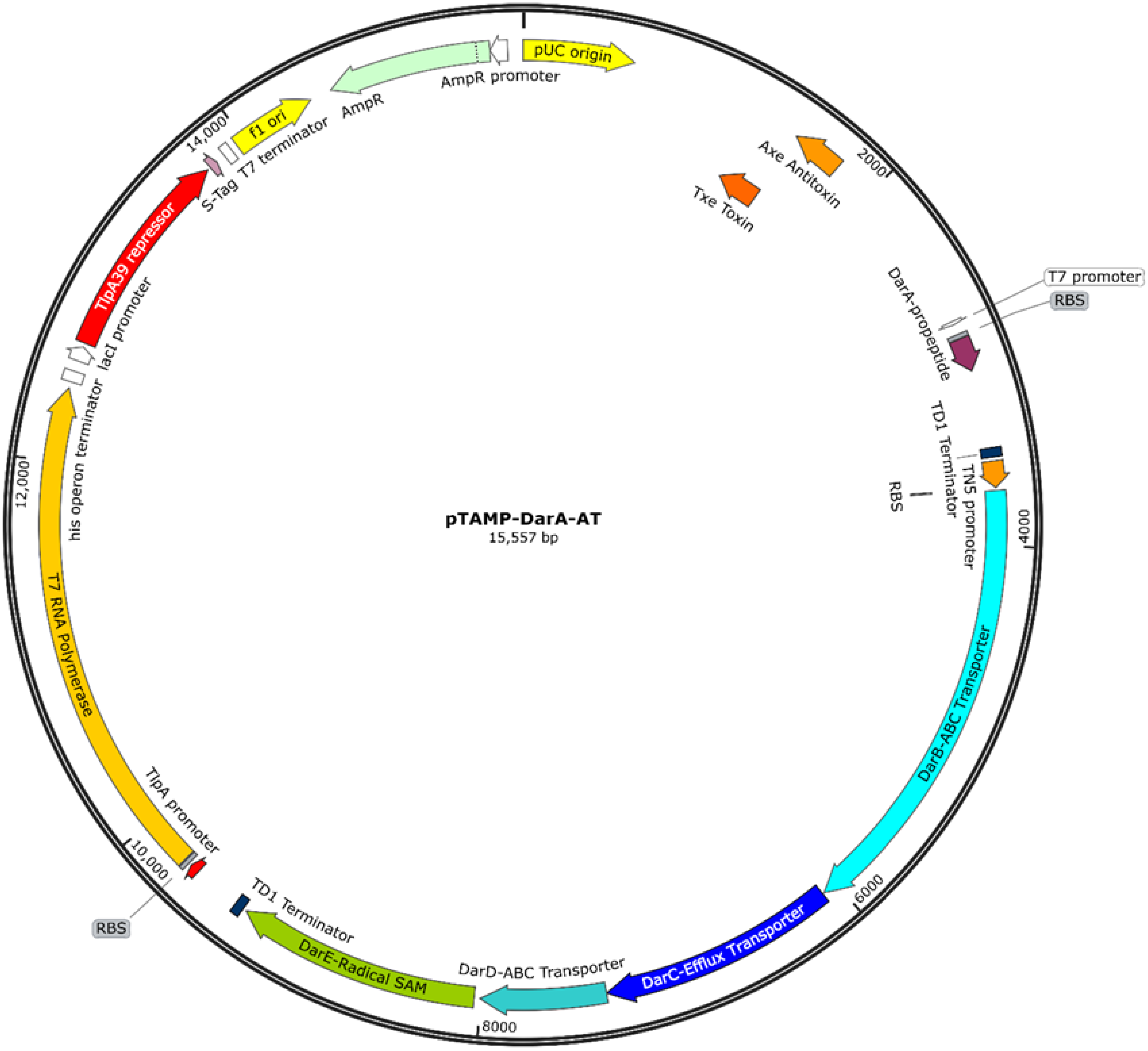


**Supplemental Figure 9.** Sequence annotated map of the pTAMP-DarA-AT recombinant plasmid

**Table S1.** Nucleotide Sequences of the genetic modules used in this study

| NAME | SEQUENCE (5' → 3') |
|---|---|
| ***darA*** | atgcataataccttaaatgaaaccgttaaaactcaagaagcactcaattctcttgctgcatcattcaaagagactgaactctcaattactgataaagcactaaacgaattaa<br>gcaataaacctaagatccctgagatcacggcctggaactggtcaaaaagcttccaggaaatttaa |
| ***darB*** | atgaatgctattaacatctcagaatccatgaaacgtttcaatatattggtatcgatactctcagtatcaatagcaatattagcaatttatttactttctactgccttctat<br>tcagatgctcgagttgataactattatcaagataatggcaacatttaccgggtggaaaccacctttaatttgcccaatggagaaaaagtgagatcggcaaaatcgcc<br>attgccgctcatagacgagctagaaaaagataaaaggataaatcgtgcagactactttttcaaacttaacacgacgattgagttacaaggaaaaaagatcaccaa<br>agttcccgtgtttgcagtcagtcatcaattcctgaataccctgtctccctttaagaaaatcgataatatcctgggagcaaatgagatttacatcacaaaagagttcaat<br>aatcgctatctcggcctggaaaatcccaaaggaaaatctattgttctggataatgaaacctatatcatcaaggatatcgtagaaaaacgtcatgattcgagtcttaat<br>atgaatgctattatttcatttaagccaacacttatcaaaaactacaataaagaaatattcaattggtatgatacacatgcttatatttttataaaactctctgatgaaa<br>aatcaatatccaatcgcgatgcgcttcttaatcatattattgcaaccaaagcccccaatcttccaggcgctccatttactgccagtgaatttattcacctatcattaaaa<br>aaatcaaaagatattcattaccaagatggattgcctgatgaaatttctattggtttatcaaaggagataatttatagctcatatatcgcgttcgcatttatcttattttcc<br>acattaaccaactattataatttaacttctgctgattcagcagaaaaaagggatatttatcagttaaagaaagcgctaggcgcttcaactttcaatattatttgcgatt<br>cactgccggccatgttgatcaaaataatcctttccacccttatctttttgttacttatcataactatatttaatgtctctctctatataaataatatatttaccctattagaa<br>gcagacagtttaatatatctagggatattttcagttgttattgcctttatctttatttcatcaatacatttttcattctcattacgtttttatatttatcaaagaaacagaaa<br>aatggatatgagatatgaatcaacccttgcctcttggtttaggaaatcatcattggctattcaattacttgtttccggaatatctatttatttagtcaccggattaaccaa<br>tcaatatattcaccttgttgatttacatactccttatgataatgccaagacaatatcaatagcaattaataacgataataataaaaaatcaactattgatgaattaagc<br>tatgagtttatcagaaaatacaatacttcaaaaataaccctcagtaattggcgaccatttgatatgtcaagagaaaatatcactattaattattccggccaacaaaa<br>caggaataactatatctcttcaaatgttattactgcggatgaaaatttcactgatgtgtggagaatgaagatattagccggtggtgataaacgagtatatcagagtg<br>ataatgctgatgtagtccatgcattagtgacaaaagaatttttaaaacagaatggaattctcaaatatgatgatatttttaacaactactattggtacacaatggagg<br>ataaaaagatccaaatcagatttattcagattatagatgatttcaatttaggtgctgtagatgaaccattcaggccaatagttgttttcatcaaaaaagatcatggaa<br>aatatgcttcgctgaatttaaacaatatgaaaaacctttctccggtacttaacgaactatcaaaacagggttttgataataatgaaattgatttaaccagtcacttatt<br>taactcttattatgataattatttaaaagtaataaaattatcatctgccgtcgctattttctcggttctattattagtcatcgcctgctttactaccagcatgactgacttta<br>acgcgatgaaaagggagttaagtataatggaatctattggtggttctatttacaccaatctcattcactttttatcgaaaagtgtcatcccgataatcgcatcaattttc<br>gtggcatatatatcaggaagaactattttagtttatctgcttgatgaatatagtgtgatttacaccgattcttttgtatacagtgctatcgccctatctgctacaatgata<br>ttcaccgttattataatgatgactacctatttggtgaattatagaaaattatccgccactaaatttttgtga |
| ***darC*** | atggatatagaaatcagaaaaaaaggcatacataacagagagaaattactcattctcattttctttatcgcttttatttcatttgttgtatttattatctattatgtcgtttcggctaa<br>agacgctttcgtgaatgaaaatgaagttacccagtttcaggtcagcagcaaagtagatagtgatattctgaaaaccagggcagttgtggttcctgataaaagtgtctctat<br>ctcaaccgagataggggggaatcgttaccgatataatgaaaaaaccatcacaagatgttttaaagaatgaagaaatcgtgaaattatctaattttaatttcacattgaataatt<br>cttcgatgttagcggacgtgacagataaactcaataatttaataaacataagaataaacttgcaatctgactatcgggatatcaataaccgattcttagaagccgccaaag<br>aactaaaagaagtagaagataaattaaagcgatatcacgatcttgtacataaaaattatatttctaaagaaactatgtccgatttaaggattaaaagagattattggcacgat<br>gtttatctattctacaaaaaattaaaaaatgataaagatcgtgatatatcaaaacagttaaaggaaatcgacgagtttgtcgaaaaacagagaaaactctctgacatcatcg<br>aaaatggtttcgagcagctttcaataaaatcccctatcgccggtaatatcagttcattagatttaattttaggacaacggttaaaacctggcgataaaatagctattgttgatg<br>atttatcaaacttttattttgaatcagaaatcaacgaatactatttaaacaaaataacataccattcatctgccagtttaatctataacaatggaaaaatacctttactggtgaaa<br>ttaatttcatctgaagtgaataatggaacattcaaagtcagatttgaattggaagataaaaaaacaattaacttcaaacgggggcaatccgttgatgtcatgattaaacttga<br>tgaagatcggaaaatattctccgtaccttcatctatggtgttttctattgacaataaaaactatgtgtttgtttaccatccagataaaaaaattgcccgacgtattcaagtttcca<br>ccggccaagataatggttctgatattgagatcaaaagtggtgttactgaaggacaaacgctcgttagctttggtaagaataaattagttaataatgatacagtaaggattg<br>aataa |
| ***darD*** | atgattagcatgatgaatatctgtaaatcatataaaacgaaattcattcaaacaaatgttttgaatgacataaatctcaatatagataaaggcgaatttatttctata<br>atgggagcgtcagggtcagggaaatcaacattactgaatgtcattggcatgtttgaaacaatcgatagcggtcaattaacactgaacaatagcaatattttaacaat<br>gaaatattctgaaaaaatatcattcagacgagaatttattggttatatttttcaatcattcaatctgttacccaatttgacggtatttgaaaacatagagttaccgttaa<br>aatacagaggttttcctaaaaaacagcggaaaagcaaagtatttgatgcaataaatagttttggattagaaaatcgggaaaatcataaaccgatacaattatctgg<br>tggacaacagcaacgtgttgctatcgccagagcaatgatagcaaaccccaccattttattagctgatgaaccaacggggaacctggatagcgtaaatggtcagaat<br>attttatcttcacttaaggagctgaatgaaaacggtacgacgattgttatggtaacccattccgtagaagcggcagcgttttctgacagaatattaacgatgagggat<br>ggccacttgcttgtttga |
| ***darE*** | atggacacaataatccccataaaatatttagattcagacgaatcatcgattcttaagaaatcatctaaaattaactacaggcaattagcttgcagaattatcggtgaaatctcc<br>gccgaaaaaatattagatgatgatgaactggctttatataataaagaaatcagtatacatttcagccctgaaattattaatgctaataaattagttgtggttgtgaaagccacc<br>aggctttgcaatttaagatgcacttattgtcactcctgggcagaaggaaaaggaaataccttaacattcttcaatttaatgcgttccattcaccgtttcttatccctaccgaata<br>tcaagcgatttgaattcgtctggcatggcggcgaagtaacgttgttgaatgttaattactttaagaaactcatctggttacaggaacaatttaaaaaaccggatcaagttatc<br>accaattcggtacagacaaatgccgtcaatattcctgaagattggttagtgttcctcaaaggtattggaatgggggtaggaataagcgttgatggtattccggaaatacac |

| | |
|---|---|
| | gatagcaggagattagattacagaggaaggccaacatcccataaagtcgcggcaagtatgaaaaagttaagaagttatggcataccttacggtgcgcttatcgtcgtcg<br>accgcgatgtttatgaatcaaatatagaaaaaatgctctcttatttttacgaaatcggtttaacggatattgaatttctgaatattgtcccagataaccgatgccagccgggtg<br>atgatcctggaggaagttatataacttaccataactatattaatttcctttctaaggttttccgtgtctggtggaatggttatcaaggcaaaatcaatattcgcttgtttgacggat<br>ttattgacagtatcaaatcgtcccaaaagaaaatgtcagattgttattgggcgggtaactgttctcaggaaataatcacattagaacctaatggtacggtatcagcatgtgat<br>aaatatgttggtgctgaagggaataattatggttcgattattgataatgatcttgggaatttactatctaaatcaaatacaaataaggatcatcttaaagaggaaatggaatctt<br>atgaaaaaatgcatcaatgtaaatggtttcatttgtgtaatggtggatgcccacacgatcgagtgaccaacaggaagcacaatccaaattatgatggttcatgttgtggaa<br>ccggcggtttgttggagacaataaaacaaaccatcgcggcgtaa |
| ***tlpA*$_{39}$**<br>**repressor** | atgcgtccggcgacatacgaaccagaacagattattgaagcagggctggccctgcaggctgaaggacggaatatcaccgggttcgcactacgtaaccaggtgggt<br>ggcggcaatccgacacgtctccgccagatatgggacgaataccaggcttcacagagcacggtcgtcactgaacccgttgccgagctgccagtggaagtggctgaa<br>gaagtgaaggccgtctccgccgcgctgtccgaacgcatcacccagctggcgacagaactgaatgacaaggcggtccgggctgcagaacgccgggttgcggaagt<br>cacgcgtgctgccggtgaacagaccgcacaggcagagcgggagctggccgacgccgcgcagacagtcgacgacctggaagaaaaactggttgaactgcaggac<br>agatatgacagtttgacgctggcgctggagtcagaacgttcactgcgtcagcagcatgatgtggagatggcccagctgaaagagcgtcttgcggccgctgaagaga<br>atacccgtcagcgagaggaacggtatcaggagcagaagacagtgctgcaggatgcgcttaatgcggagcaggcacagcacaaaaacacgcgggaagacctgca<br>gaaacgactggagcaaatttctgtcgaagctaatgcgcgtacagaagaactgaagtctgaacgcgataaagtcaatactttccttacccgccttgaatcgcaggaa<br>aatgcgctggcctcagaacgtcagcagcatctggccacccgcgaaacgctgcagcaacgcctcgagcaggccatcgctgacacgcaggcgcgcgccggtgagatt<br>gcacttgaacgtgacagagtcagcagcctcaccgcaaggctggaatcgcaggaaaaggcctcctcggagcaactggtgcgtatgggcagtgaaatagccagtctg<br>acagagcgttgcacacagctggaaaaccagcgtgatgatgcccgtctggagacgatgggggagaaagaaacggtcgcggcactgcgtggtgaggctgaagccct<br>gaagcgtcagaaccagtcactgatggcggcgctttcaggcaataaacagaccggtggccagaatgcgtga |
| ***txe***<br>**toxin** | atgattaaggcttggtctgatgatgcttgggatgattatctttattggcatgagcaaggaaacaaaagcaatataaaaaagattaacaagttaataaaagatatcg<br>atcgttccccctttgctggattaggaaaacctgagccattaaagcatgatttatctggaaaatggtccagaagaattacagatgaacatagactgatatatagagtt<br>gaaaatgaaacgatatttatttattctgcaaaagatcactattaa |
| ***axe***<br>**antitoxin** | atggaagcagtagcttattcaaatttccgccaaaatttacgtagttatatgaaacaagttaatgaggatgctgaaacacttattgtaacaagtaaagatgtagaaga<br>tacagttgttgtattatcaaaaagagattatgattctatgcaagaaacgttgagaacactttctaataattacgtcatggaaaaaattcgtcgaggagatgaacaatt<br>ctccaaaggtgcatttaaaacacatgacttaatcgaggttgaatctgatgattaa |
| ***t7***<br>***RNAP*** | atgaacacgattaacatcgctaagaacgacttctctgacatcgaactggctgctatcccgttcaacactctggctgaccattacggtgagcgtttagctcgcgaacag<br>ttggcccttgagcatgagtcttacgagatgggtgaagcacgcttccgcaagatgtttgagcgtcaacttaaagctggtgaggttgcggataacgctgccgccaagcc<br>tctcatcactaccctactccctaagatgattgcacgcatcaacgactggtttgaggaagtgaaagctaagcgcggcaagcgcccgacagccttccagttcctgcaag<br>aaatcaagccggaagccgtagcgtacatcaccattaagaccactctggcttgcctaaccagtgctgacaatacaaccgttcaggctgtagcaagcgcaatcggtcg<br>ggccattgaggacgaggctcgcttcggtcgtatccgtgaccttgaagctaagcacttcaagaaaaacgttgaggaacaactcaacaagcgcgtagggcacgtctac<br>aagaaagcatttatgcaagttgtcgaggctgacatgctctctaagggtctactcggtggcgaggcgtggtcttcgtggcataaggaagactctattcatgtaggagt<br>acgctgcatcgagatgctcattgagtcaaccggaatggttagcttacaccgccaaaatgctggcgtagtaggtcaagactctgagactatcgaactcgcacctgaat<br>acgctgaggctatcgcaacccgtgcaggtgcgctggctggcatctctccgatgttccaaccttgcgtagttcctcctaagccgtggactggcattactggtggtggcta<br>ttgggctaacggtcgtcgtcctctggcgctggtgcgtactcacagtaagaaagcactgatgcgctacgaagacgtttacatgcctgaggtgtacaaagcgattaaca<br>ttgcgcaaaacaccgcatggaaaatcaacaagaaagtcctagcggtcgccaacgtaatcaccaagtggaagcattgtccggtcgaggacatccctgcgattgagc<br>gtgaagaactcccgatgaaaccggaagacatcgacatgaatcctgaggctctcaccgcgtggaaacgtgctgccgctgctgtgtaccgcaaggacaaggctcgca<br>agtctcgccgtatcagccttgagttcatgcttgagcaagccaataagtttgctaaccataaggccatctggttcccttacaacatggactggcgcggtcgtgtttacgc<br>tgtgtcaatgttcaacccgcaaggtaacgatatgaccaaaggactgcttacgctggcgaaaggtaaaccaatcggtaaggaaggttactactggctgaaaatccac<br>ggtgcaaactgtgcgggtgtcgataaggttccgttccctgagcgcatcaagttcattgaggaaaaccacgagaacatcatggcttgcgctaagtctccactggagaa<br>cacttggtgggctgagcaagattctccgttctgcttccttgcgttctgctttgagtacgctggggtacagcaccacggcctgagctataactgctcccttccgctggcgt<br>ttgacgggtcttgctctggcatccagcacttctccgcgatgctccgagatgaggtaggtggtcgcgcggttaacttgcttcctagtgaaaccgttcaggacatctacgg<br>gattgttgctaagaaagtcaacgagattctacaagcagacgcaatcaatgggaccgataacgaagtagttaccgtgaccgatgagaacactggtgaaatctctga<br>gaaagtcaagctgggcactaaggcactggctggtcaatggctggcttacggtgttactcgcagtgtgactaagcgttcagtcatgacgctggcttacgggtccaaag<br>agttcggcttccgtcaacaagtgctggaagataccattcagccagctattgattccggcaagggtctgatgttcactcagccgaatcaggctgctggatacatggcta<br>agctgatttgggaatctgtgagcgtgacggtggtagctgcggttgaagcaatgaactggcttaagtctgctgctaagctgctggctgctgaggtcaaagataagaag<br>actggagagattcttcgcaagcgttgcgctgtgcattgggtaactcctgatggtttccctgtgtggcaggaatacaagaagcctattcagacgcgcttgaacctgatg<br>ttcctcggtcagttccgcttacagcctaccattaacaccaacaaagatagcgagattgatgcacacaaacaggagtctggtatcgctcctaactttgtacacagcca<br>agacggtagccaccttcgtaagactgtagtgtgggcacacgagaagtacggaatcgaatcttttgcactgattcacgactccttcggtaccattccggctgacgctgc<br>gaacctgttcaaagcagtgcgcgaaactatggttgacacatatgagtcttgtgatgtactggctgatttctacgaccagttcgctgaccagttgcacgagtctcaatt<br>ggacaaaatgccagcacttccggctaaaggtaacttgaacctccgtgacatcttagagtcggacttcgcgttcgcgtaa |
| | atgcaagcggcaactgttgtgattaaccgccgcgctctgcgacacaacctgcaacgtcttcgtgaactggcccctgccagtaaaatggttgcggtggtgaaagcga<br>acgcttatggtcacggtcttcttgagaccgcgcgaacgctccccgatgctgacgcctttggcgtagcccgtctcgaagaagctctgcgactgcgtgcgggggggaatc<br>accaaacctgtactgttactcgaaggctttttgatgccagagatctgccgacgatttctgcgcaacattttcataccgccgtgcataacgaagaacagctggctgcg |

| | |
|---|---|
| **alanine racemase (*alr*)** | ctggaagaggctagcctggacgagccggttaccgtctggatgaaactcgataccggtatgcaccgtctgggcgtaaggccggaacaggctgaggcgttttatcatc<br>gcctgacccagtgcaaaaacgttcgtcagccggtgaatatcgtcagccattttgcgcgcggatgaaccaaaatgtggcgcaaccgagaaacaactcgctatctt<br>taataccttttgcgaaggcaaacctggtcaacgttccattgccgcgtcgggtggcattctgctgtggccacagtcgcattttgactgggtgcgcccgggcatcattcttt<br>atggcgtctcgccgctggaagatcgctccaccggtgccgattttggctgtcagccagtgatgtcactaacctccagcctgattgccgtgcgtgagcataaagccgga<br>gagcctgttggttatggtggaacctgggtaagcgaacgtgatacccgtcttggcgtagtcgcgatgggctatggcgatggttatccgcgcgccgcgccgtccggtac<br>gccagtgctggtgaacggtcgcgaagtaccgattgtcgggcgcgtggcgatggatatgatctgcgtagacttaggtccacaggcgcaggacaaagccggggatcc<br>ggtcattttatggggcgaaggtttgcccgtagaacgtatcgctgaaatgacgaaagtaagcgcttacgaacttattacgcgcctgacttcaagggtcgcgatgaaat<br>acgtggat |

**Table S2.** List of primers and their corresponding sequences used for qRT-PCR and qPCR analysis

| Primer Name | Sequence (5'→ 3') | Description | Reference |
|---|---|---|---|
| **darA Fwd** | GCACTCAATTCTCTTGCTGCAT | qRT-PCR forward primer for darA | This study |
| **darA Rev** | TTTTGACCAGTTCCAGGCCG | qRT-PCR reverse primer for darA | This study |
| **T7RNAP Fwd** | CGAGAACATCATGGCTTGCG | qRT-PCR forward primer for T7RNAP | This study |
| **T7RNAP Rev** | CCCAGCGTACTCAAAGCAGA | qRT-PCR reverse primer for T7RNAP | This study |
| **pUC Fwd** | TTGCCGGATCAAGAGCTACC | qPCR forward primer for pUC origin | This study |
| **pUC Rev** | GGCGGTGCTACAGAGTTCTT | qPCR reverse primer for pUC origin | This study |
| **16S rRNA Fwd** | CATGCCGCGTGTATGAAGAA | qRT-PCR forward primer for 16S rRNA | Smati et al., 2013 |
| **16S rRNA Rev** | CGGGTAACGTCAATGAGCAAA | qRT-PCR reverse primer for 16S rRNA | Smati et al., 2013 |
